# Preclinical validation of a repurposed metal chelator as a community-based therapeutic for hemotoxic snakebite

**DOI:** 10.1101/717280

**Authors:** Laura-Oana Albulescu, Melissa Hale, Stuart Ainsworth, Jaffer Alsolaiss, Edouard Crittenden, Juan J. Calvete, Mark C. Wilkinson, Robert A. Harrison, Jeroen Kool, Nicholas R. Casewell

## Abstract

Snakebite envenoming causes 138,000 deaths annually and ~400,000 victims are left with permanent disabilities. Envenoming by saw-scaled vipers (Viperidae: *Echis*) leads to systemic hemorrhage and coagulopathy, and represents a major cause of snakebite mortality and morbidity in Africa and Asia. The only specific treatment for snakebite, antivenom, has poor specificity, low affordability, and must be administered in clinical settings due to its intravenous delivery and high rates of adverse reactions. This requirement results in major treatment delays in resource-poor regions and impacts substantially on patient outcomes following envenoming. Here we investigated the value of metal chelators as novel community-based therapeutics for snakebite. Among the tested chelators, dimercaprol (British anti-Lewisite) and its derivative 2,3-dimercapto-1-propanesulfonic acid (DMPS), were found to potently antagonize the activity of Zn^2+^-dependent snake venom metalloproteinase toxins *in vitro*. Moreover, DMPS prolonged or conferred complete survival in murine preclinical models of envenoming against a variety of saw-scaled viper venoms. DMPS also significantly extended survival in a ‘challenge and treat’ model, where drug administration was delayed post-venom injection, and the oral administration of this chelator provided partial protection against envenoming. Finally, the potential clinical scenario of early oral DMPS therapy combined with a later, delayed, intravenous dose of conventional antivenom provided prolonged protection against the lethal effects of envenoming *in vivo*. Our findings demonstrate that safe and affordable repurposed metal chelators effectively neutralize saw-scaled viper venoms *in vitro* and *in vivo* and highlight the great promise of DMPS as a novel, community-based, early therapeutic intervention for hemotoxic snakebite envenoming.

## Introduction

Snakebite envenoming is a neglected tropical disease (NTD) that affects ~5 million people worldwide each year, and leads to high mortality (~138,000/year) and morbidity (~400,000-500,000/year), particularly in sub-Saharan Africa and southern Asia *(1)*. In West Africa, the economic burden of snakebite has been estimated at 320,000 Disability Adjusted Life Years (DALYs) annually - a number higher than most other NTDs, including leishmaniasis, trypanosomiasis and onchocerciasis *(2)*. The highest burden of snakebite is suffered by the rural impoverished communities of low/middle income countries, who often rely on agricultural activities for their income *(3)*. These activities put them at risk of snakebite by being exposed to environments inhabited by venomous snakes, and the remoteness of many of these communities makes accessing healthcare problematic *(4)*. Consequently, it is estimated that 75% of snakebite fatalities occur outside of a hospital setting *(5)*, as victims are often delayed in reaching a healthcare facility due to long travel times and/or suboptimal health seeking behaviors. Crucially, treatment delays are known to result in poor patient outcomes, and often lead to life-long disabilities, psychological sequelae or death *(6, 7)*. Further compounding this situation, species-specific antivenom, the only appropriate treatment for snakebite *(1, 3)*, is often unavailable locally and, when present, is exceedingly expensive relative to the income of snakebite victims, despite being classified by the WHO as an essential medicine *(8)*. Consequently, snakebite was classified as a ‘priority NTD’ by the World Health Organization (WHO) in 2017. Subsequently, the WHO has developed a strategy proposed to halve the number of snakebite deaths and disabilities by the year 2030, by improving existing treatments, developing novel therapeutics, and empowering local communities to improve prehospital treatment *(9)*.

Snake venoms are mixtures of numerous toxin isoforms encoded by multiple gene families, whose composition varies both inter- and intra-specifically *(10, 11)*. Venom variation makes the development of pan-specific snakebite treatments challenging due to the multitude of drug targets that need to be neutralized for any particular geographical region. Antivenoms consist of polyclonal antibodies (IgG or IgG-derived F(ab’)_2_ or Fab fragments) purified from the serum/plasma of animals hyperimmunized with snake venom/s. Venom toxin variation results in these products typically having limited or no efficacy against snake venoms not used in the manufacturing process *(12)*. Furthermore, antivenoms have poor dose efficacy (only ~10-20% of antibodies are specific to venom toxins *(13)*), must be delivered intravenously, and have a high incidence of adverse reactions (as high as 75% of cases *(14)*), meaning that they must be delivered in a clinical setting *(1)*. As advocated for by the WHO *(9)*, there is an urgent need for the development of rapid, effective, well-tolerated and affordable interventions for treating tropical snakebite *(9)*. Ideally, initial interventions would be safe for use outside of a clinical environment, thus facilitating rapid administration in a community setting and likely improving patient outcomes.

The use of small molecule inhibitors to generically target key classes of snake venom toxins has recently attracted renewed interest as a potential therapeutic alternative to antivenom *(15)*. Varespladib, a secretory phospholipase A2 (PLA_2_) inhibitor, has emerged as a promising drug candidate that protects against venom-induced lethality and limits the pathology associated with certain snakebites in preclinical animal models *(16, 17)*. Varespladib was found to be particularly effective against PLA_2_-rich elapid venoms, but also showed some efficacy against a number of viper venoms *(18)*. Molecules that counteract the activity of snake venom metalloproteinase (SVMPs) toxins have also been investigated in this regard. SVMPs are Zn^2+^-dependent, enzymatically active hemotoxins *(19)* involved in causing systemic hemorrhage and coagulopathy by actively degrading capillary basement membranes and/or interacting with key components of the blood clotting cascade *(20)*. They are often the major toxin constituents of viper venoms *(21)*. Peptidomimetic hydroxamate inhibitors that block the catalytic site of SVMPs (e.g. batimastat, marimastat) have been shown to abolish the local and systemic toxicity induced by saw-scaled viper (*Echis ocellatus*) venoms from Cameroon and Ghana *(22)*, and dermonecrosis and hemorrhage caused by the venom of the Central American fer-de-lance (*Bothrops asper*) *(23)*. Similarly, metal chelators that deplete the Zn^2+^ pool required for SVMP function, including EDTA *(23, 24)*, can prevent hemorrhage, myotoxicity and/or lethality caused by *E. ocellatus* or *E. carinatus* venoms *(25, 26)*. These previous studies suggest that small molecule inhibitors targeting SVMPs may represent good candidates for delaying or even neutralizing the toxicity caused by SVMP-rich venoms. However, neither marimastat nor batimastat are currently available as licensed medicines, whereas EDTA is not desirable for further development as a therapeutic due to its poor specificity for zinc and high affinity for calcium, its poor safety profile, as well as a requirement for slow intravenous administration, which makes it unsuitable for use as a community therapeutic *(27)*.

Saw-scaled vipers (Viperidae: *Echis* spp.) represent one of the most medically-important groups of venomous snakes, with a broad distribution through Africa north of the equator, the Middle East and western parts of south Asia (Fig. 1). Among these, *E. ocellatus* (the West African saw-scaled viper) is responsible for most snakebite deaths in sub-Saharan Africa *(28)*, with a case fatality rate of 20% in the absence of antivenom treatment *(29)*, while the Indian saw-scaled viper (*E. carinatus*) is one of the ‘big four’ snake species that together cause the vast majority of India’s 46,000 annual snakebite deaths *(5)*. Envenomings by saw-scaled vipers cause local tissue damage and venom-induced consumption coagulopathy (VICC), the combination of which routinely leads to life-threatening internal hemorrhage *(30, 31)*. Crucially, the predominant toxins in saw-scaled viper venoms are typically SVMPs (27.4-75.7% of all venom proteins; Fig. 1) *(11, 24)*, and thus these snakes are tractable models for testing the development of new SVMP-specific small molecule-based snakebite therapeutics. Consequently, in this study we investigated the *in vitro* and *in vivo* neutralizing potential of repurposed, licensed, metal chelators against a variety of saw-scaled viper venoms (genus *Echis*). We demonstrate that 2,3-dimercapto-1-propanesulfonic acid (DMPS), an existing licensed drug indicated for treating chronic and acute heavy metal poisoning, effectively neutralizes venom-induced lethality *in vivo* and is a highly promising candidate for translation into a new community-based treatment for snakebite envenoming.

**Fig. 1.**
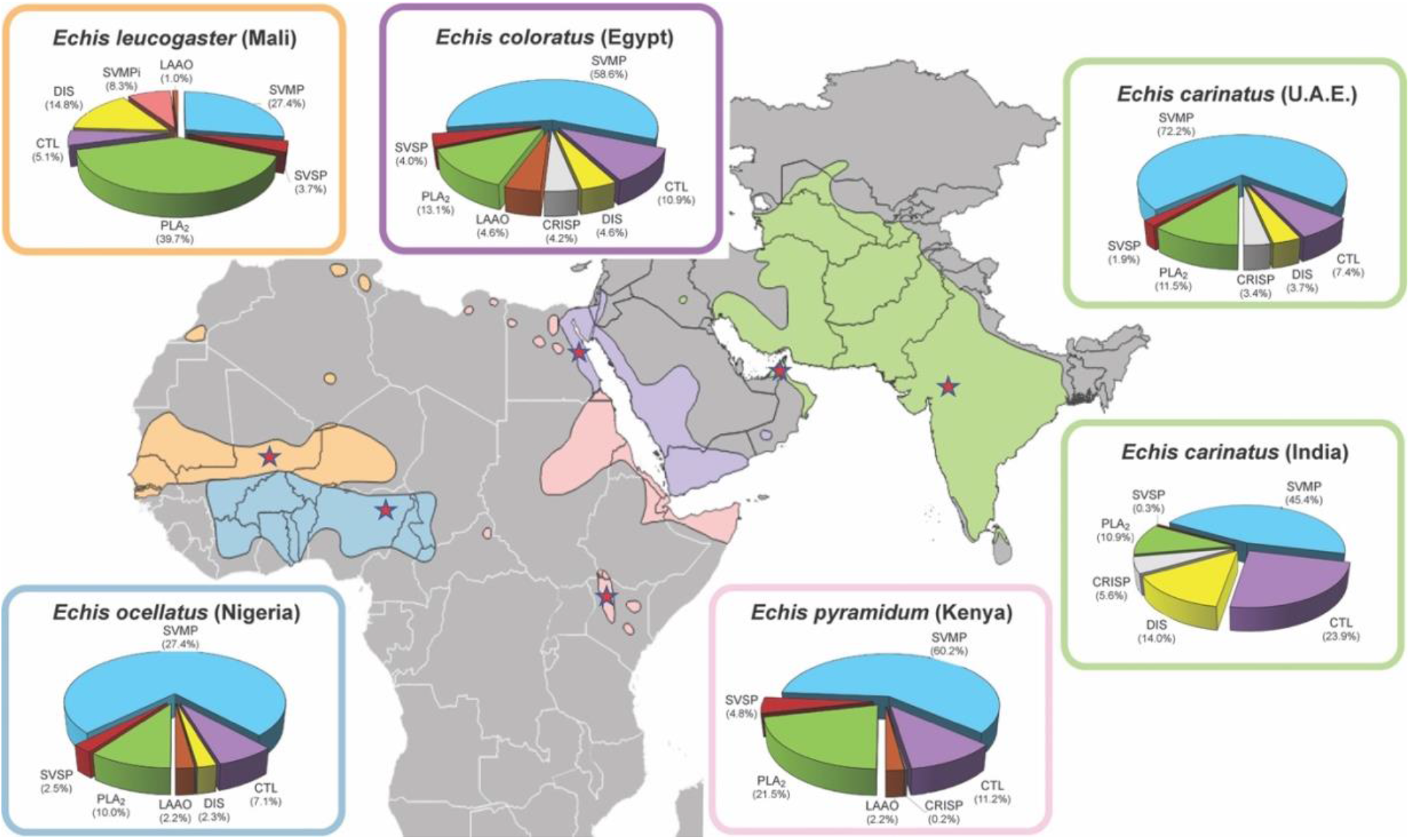
The geographical distribution and venom toxin composition of saw-scaled vipers (genus *Echis*) Map of saw-scaled viper distribution with the locales of the studied species indicated by red stars. *Echis* distribution areas and corresponding venom proteomes are highlighted by the following colors: light orange (*E. leucogaster*), blue (*E. ocellatus (11)*), green (*E. carinatus*), pink (*E. pyramidum (11)*), violet (*E. coloratus (11)* Toxin proteome abundances were taken from *(11, 57)* and generated in this study for *E. leucogaster*. Toxin family key: SVMP, snake venom metalloproteinase; SVSP, snake venom serine proteinase; PLA2, phospholipase A2; CTL, C-type lectins; LAAO, L-amino acid oxidase; SVMPi, SVMP inhibitors; DIS, disintegrin; CRISP, Cysteine-rich secretory protein.

## Results

### Metal chelators inhibit venom activity in vitro

Saw-scaled viper venoms are highly enriched in SVMPs (45.4-72.5% of all toxins), with the exception of *E. leucogaster*, whose venom proteome is dominated by PLA_2s_ (39.7%; SVMPs 27.4%) (Fig. 1). As SVMP activity is Zn^2+^-dependent, we assayed the ability of three metal chelators to inhibit venom SVMPs from six saw-scaled vipers with a wide geographical distribution (Fig. 1), namely *E. ocellatus* (Nigeria), *E. carinatus* (U.A.E. and India), *E. pyramidum* (Kenya), *E. leucogaster* (Mali), and *E. coloratus* (Egypt). The three chosen chelators - dimercaprol (British anti-Lewisite), 2,3-dimercapto-1-propanesulfonic acid (DMPS, also known as unithiol), and dimercaptosuccinic acid (DMSA, also known as succimer) – share a common backbone and are licensed medicines used to treat acute and chronic heavy metal poisoning *(32)*. We also compared the efficacy of these chelators to that of EDTA, a metal chelator that we have previously shown to inhibit the SVMP activity and lethal effects of *E. ocellatus* venom *(24)*.

We used a kinetic chromogenic assay to quantify venom SVMP activity and its neutralization by metal chelators. All six *Echis* venoms displayed similar levels of SVMP activity (Fig. S1A), with slightly higher levels observed for *E. carinatus* (India). When testing the neutralization of this activity across a 1000-fold concentration range for the various metal chelators (150 μM to 150 nM), dimercaprol was found to consistently be the most effective (IC_50_: 0.04-0.18 μM), followed by DMPS~EDTA (IC_50_: 1.64-3.40 and 2.25-4.43 μM, respectively) and then DMSA (IC_50_=19.29-35.09 μM) (Fig. 2). However, all drugs inhibited >98% of venom activity at the maximal concentration tested (150 μM), with dimercaprol, DMPS and EDTA displaying comparable potency at the 15 μM concentration, irrespective of the snake species tested.

**Fig. 2.**
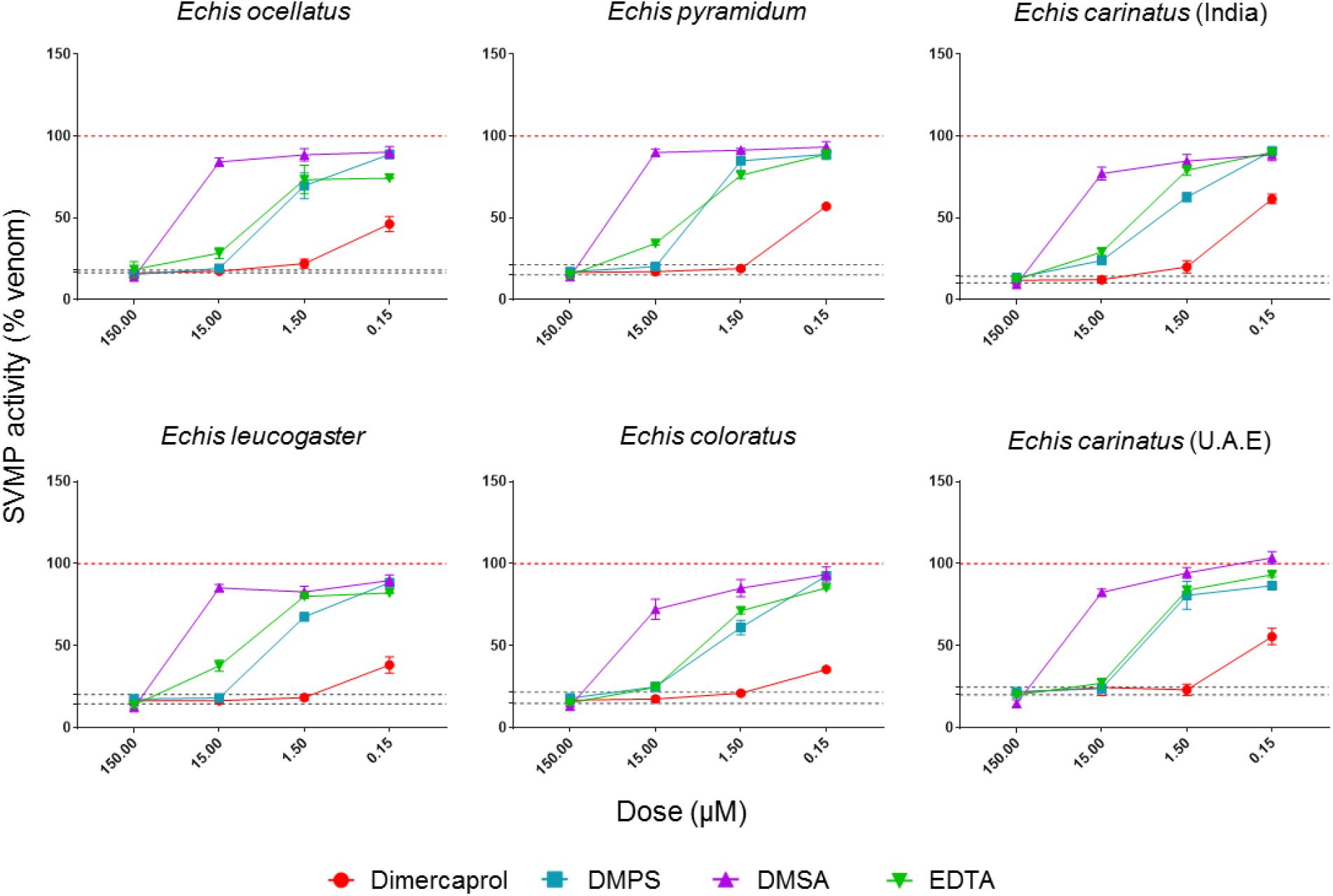
Metal chelators inhibit the snake venom metalloproteinase (SVMP) toxin activity of saw-scaled viper venoms. The neutralizing capability of four metal chelators against the SVMP activity of six *Echis* venoms. Data is presented for four drug concentrations, from 150 μM to 150 nM (highest to lowest dose), expressed as percentages of the venom-only sample (100%, dotted red line). The negative control is presented as an interval (dotted black lines) and represents the values recorded in the PBS-only samples (expressed as % of venom activity), where the highest and the lowest values for each set of experiments are depicted. Inhibitors are color-coded (dimercaprol, red; DMPS, blue; DMSA, purple; EDTA, green). The data represents triplicate independent repeats with SEMs, where each repeat represents the average of n≥2 technical replicates.

We next tested the inhibitory capacity of the chelators in a venom-induced plasma clotting assay, an *in vitro* model for *in vivo* coagulopathy *(33)*. As all tested *Echis* venoms are procoagulant *(24, 34)* and exhibit similar, rapid, clotting profiles (Fig. S1B), we measured venom neutralization as a shift in these profiles towards that of the plasma-only control. Similar to the SVMP assay, dimercaprol outperformed DMPS and DMSA (IC_50_: 2.54-32.03, 49.34-502.40 and 52.84-2042.00 μM, respectively). However, with few exceptions, all three drugs showed considerable neutralization (47.3-95.7% inhibition) of the coagulopathic effects of each of the venoms at the maximal inhibitor dose tested (150 μM) (Fig. 3).

**Fig. 3.**
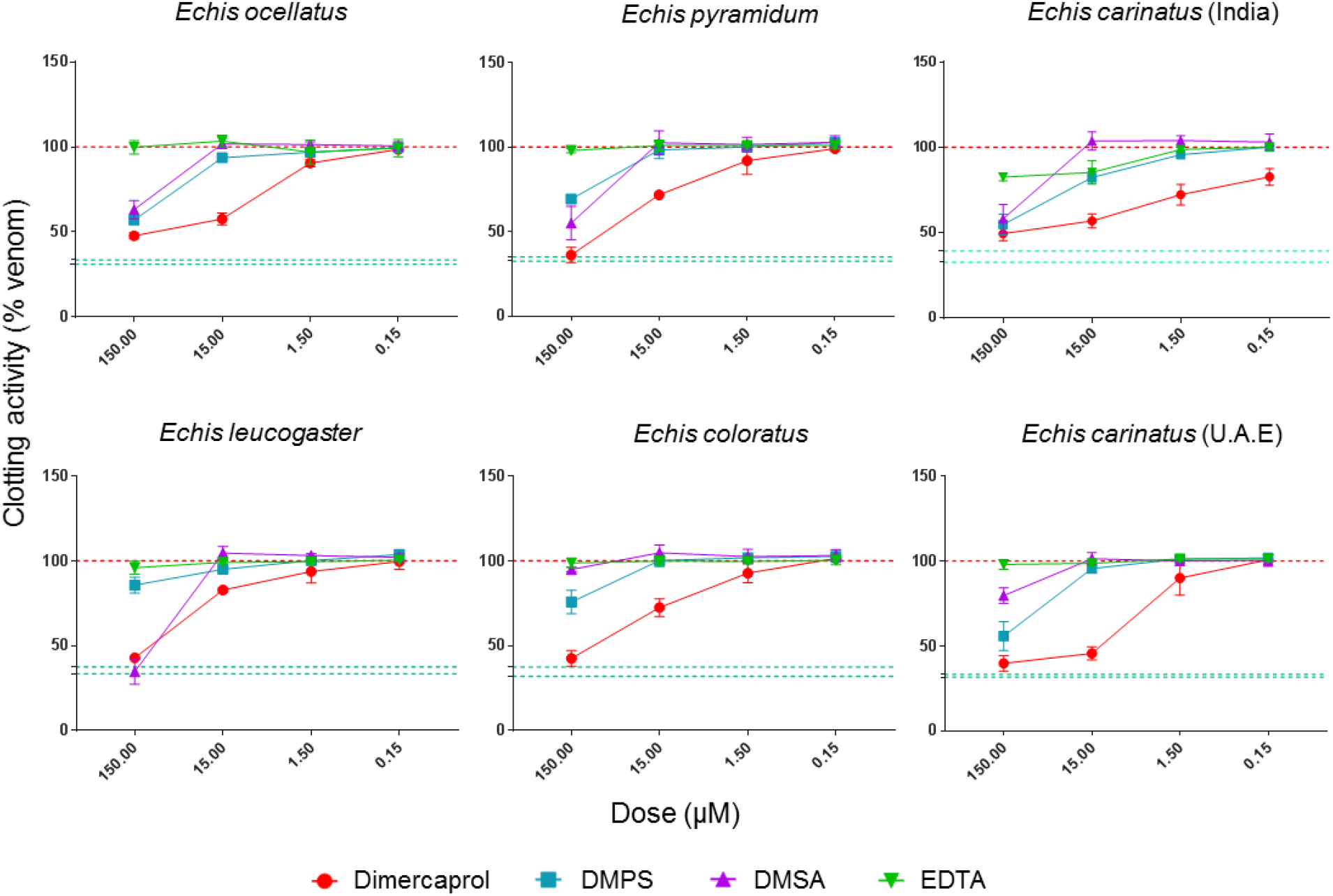
Metal chelators inhibit the procoagulant activity of saw-scaled viper venoms. The neutralizing capability of four metal chelators against the procoagulant activity of six *Echis* venoms. Data is presented for four drug concentrations, from 150 μM to 150 nM (highest to lowest dose), expressed as percentages of the venom-only sample (100%, dotted red line). The negative control is presented as an interval (dotted green lines) and represents normal plasma clotting (expressed as % of venom activity). Inhibitors are color-coded (dimercaprol, red; DMPS, blue; DMSA, purple; EDTA, green). The data represents triplicate independent repeats with SEMs, where each repeat represents the average of n≥2 technical replicates.

EDTA did not inhibit venom activity in the plasma assay due to its proclivity towards calcium, which is added to the citrated plasma in excess as a required cofactor to stimulate clotting. As five of the six *Echis* venoms are capable of inducing clotting in a calcium-independent manner *(24, 34)* (Fig. S2), we repeated the assay in the absence of calcium and in this scenario, EDTA was effective at inhibiting venom-induced plasma clotting, presumably through effective chelation of Zn^2+^ in the absence of calcium. Despite this, EDTA was generally outperformed by both dimercaprol and DMPS under these conditions (IC_50_: 1.40-261.40 versus 0.70-6.12 and 3.29-22.07 μM, respectively).

Saw-scaled viper venoms contain SVMPs that potently activate prothrombin, and therefore contribute to VICC in envenomed patients *(24)*. Given the success of metal chelators in neutralizing procoagulant venom activity in the above assays, we next tested whether they could prevent the cleavage of prothrombin. Recombinant human prothrombin was either preincubated with venom or venom and inhibitor (150 or 500 μM doses of each metal chelator) (Fig. S3), with prothrombin profiles subsequently examined by SDS-PAGE. All four chelators demonstrated protective effects against each saw-scaled viper venom, with the exception of *E. coloratus*, at the higher dose, although dimercaprol and EDTA outperformed DMPS and DMSA at the lower dose (150 μM) (Fig. S3).

### Dimercaprol and DMPS protect against venom-induced lethality

Given the *in vitro* efficacy of the tested chelators, which generally outperformed EDTA, a drug shown to neutralize the lethal effects of saw-scaled viper venoms *in vivo (24)*, we next investigated whether these compounds could protect against venom-induced lethality in a conventional *in vivo* model of envenoming. To this end, we used a refined version of the WHO-recommended standard protocol for the preclinical testing of antivenoms (ED_50_ assay). This assay requires the preincubation of venom and therapy prior to their intravenous co-administration to mice. Groups of male CD1 mice (n=5) were injected via the tail vein with 2.5 x lethal dose 50 (LD_50_) of *E. ocellatus* (Nigeria, 45 μg) venom in the presence or absence of the various chelators (60 μg dose of dimercaprol, DMPS or DMSA per mouse). We used *E. ocellatus* as our model, due to this species likely being the most medically-important snake in Africa. The positive control venom-only group all succumbed to the lethal effects of the venom within the first hour (3-50 min). However, the groups of mice that received dimercaprol or DMPS alongside a lethal dose of venom survived until the end of the experiment (6 h) (Fig. 4). DMSA was found to be less effective, as only three of the five experimental animals survived (Fig. 4C). The metal chelators were also well tolerated when administered alone, as the mice exhibited no overt signs of toxicity (e.g. normal behavior) (Fig. 4).

**Fig. 4.**
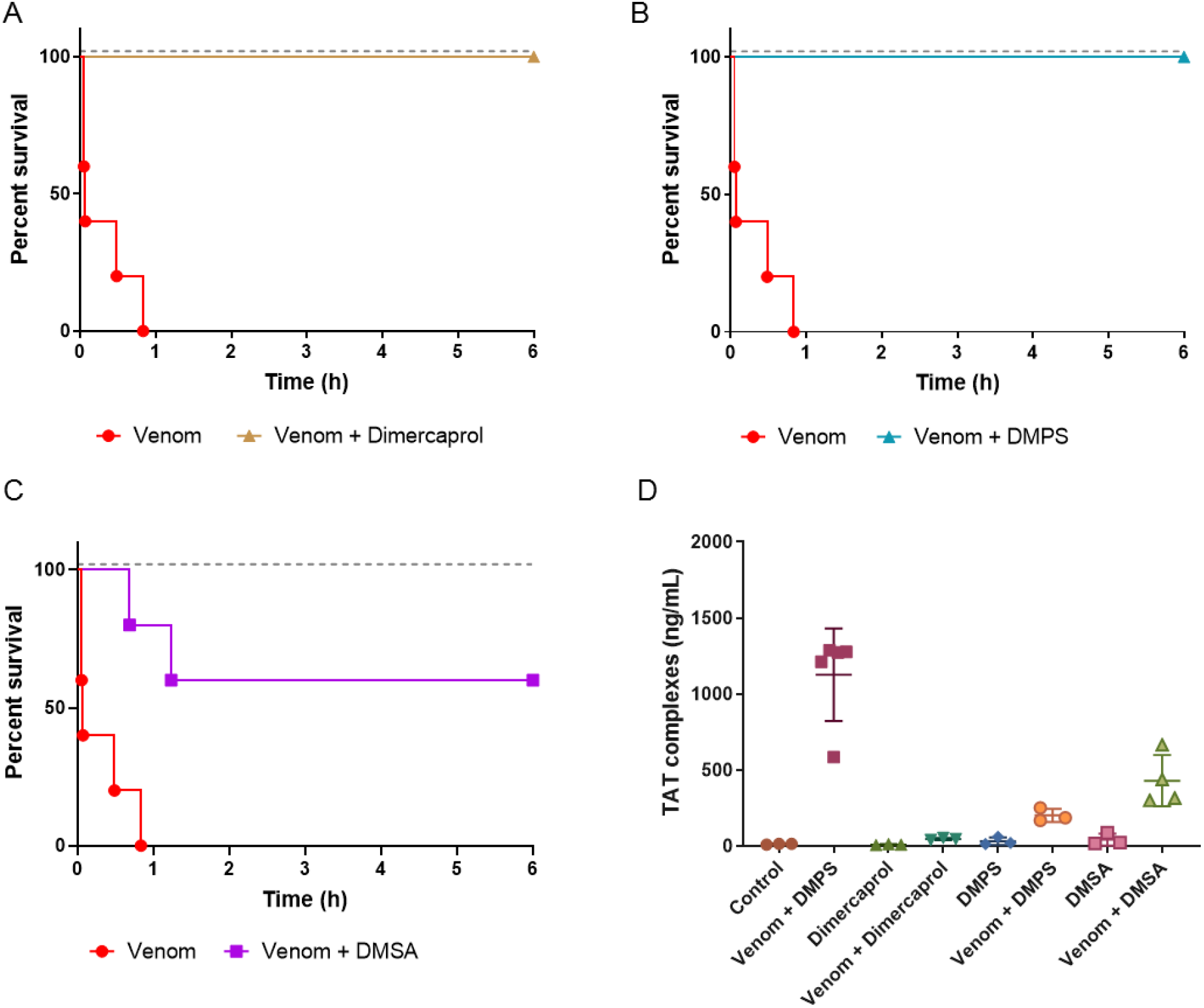
Metal chelators prevent or delay lethality *in vivo* when preincubated with *Echis ocellatus* venom. Kaplan-Meier survival graphs for experimental animals (n=5) receiving venom preincubated (30 mins at 37°C) with different metal chelators via the intravenous route. Survival of mice receiving 45 μg of *E. ocellatus* venom (2.5 × LD_50_ dose) with and without 60 μg of dimercaprol (**A**), or 60 μg of DMPS (**B**), or 60 μg of DMSA (**C**). Drug-only controls are presented as black dotted lines at the top of each graph (none of the drugs exhibited toxicity at the given doses) and the end of the experiment was at 6 h. (**D**) Quantification of thrombin-antithrombin (TAT) levels in envenomed animals. Where the time of death was the same within experimental groups (e.g. early deaths or complete survival) TAT levels were quantified for n=3, and where times of death varied n=5. The data displayed represents means of the duplicate technical repeats plus SDs.

We have previously demonstrated that plasma thrombin-antithrombin (TAT) levels, a marker of thrombin generation, represent a useful biomarker for assessing the efficacy of therapeutics against saw-scaled viper envenoming in experimental animals *(24)*. In the current study, mortality correlated with high TAT levels (~1200 ng/ml, R^2^=0.866, Fig. S4), whereas low TAT levels (Fig. 4D) were observed in mice where metal-chelators prevented lethality, and these levels were found to be comparable to those quantified from non-envenomed control animals (10.2-15.9 ng/ml). In line with our *in vitro* findings, dimercaprol caused the greatest decrease in TAT levels (~40-50 ng/ml), followed by DMPS (~170-252 ng/ml) and then DMSA (~300-666 ng/ml).

### Chelator efficacy varies among snake species

We next tested the capability of the two metal chelators that provided complete protection against the lethal effects of *E. ocellatus* envenoming (dimercaprol and DMPS) at neutralizing venom from related saw-scaled viper species. We chose the next two most medically important *Echis* species (*E. pyramidum* [Kenya] and *E. carinatus* [India]), each of which has a relatively different venom composition when compared to that of *E. ocellatus* (Fig.1, *(11, 35)*). Whilst both *E. pyramidum* and *E. carinatus* venoms remain dominated by SVMPs, they each have higher proportions of C-type lectin toxins (CTLs) (11.2% and 23.9%) and PLA_2s_ (*E. pyramidum*, 21.5%) or disintegrins (*E. carinatus*, 14.0%) than *E. ocellatus* (CTL, 7.1%; PLA_2_, 10.0%; disintegrins, 2.3%). Neither chelator was found to confer complete protection against both of these venoms when tested *in vivo*. Dimercaprol provided complete protection against lethality caused by *E. carinatus*, but was less effective against *E. pyramidum* (two deaths; 54 and 165 min) (Fig. 5A), while DMPS provided complete protection against *E. pyramidum*, but only partial neutralization against *E. carinatus* (two deaths; 89 and 110 min) (Fig. 5B). TAT levels showed a consistent correlation with the survival data, with lethality corresponding to increased TAT levels (>322 ng/ml). We find it highly promising that both metal chelators were capable of preventing lethality in mice envenomed with two of the three venoms tested, and that for the third partial protection and significantly prolonged survival times were observed (P=0.0041 for *E. carinatus* venom-only vs. venom + DMPS; P=0.0037 for *E. pyramidum* venom-only vs. venom + dimercaprol) (Figs. 4 and 5).

**Fig. 5.**
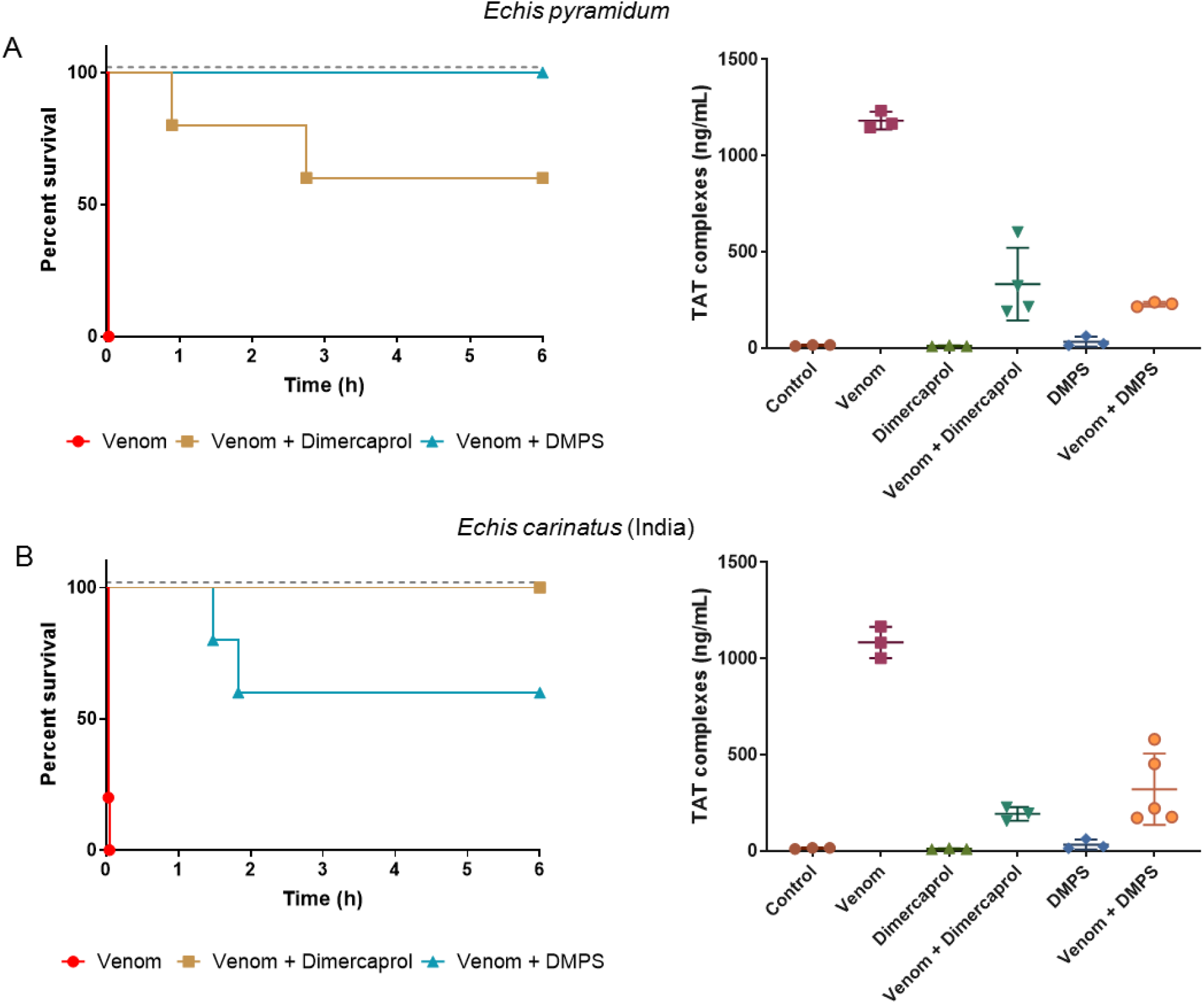
Chelator efficacy against other medically important *Echis* venoms. Kaplan-Meier survival graphs for experimental animals (n=5) receiving *Echis pyramidum and E. carinatus* (India) venom preincubated (30 mins at 37 °C) with different metal chelators via the intravenous route. Survival of mice receiving 40 μg of *E. pyramidum* venom (2.5 × LD_50_ dose) (**A**), and 47.5 μg of *E. carinatus* (India) venom (2.5 × LD_50_ dose) (**B**) with or without 60 μg of dimercaprol or DMPS. Drug-only controls are presented as black dotted lines at the top of each graph (none of the drugs exhibited toxicity at the given doses) and the end of the experiment was at 6 h. Quantification of thrombin-antithrombin (TAT) levels in envenomed animals are displayed in the right panels. Where the time of death was the same within experimental groups (e.g. early deaths or complete survival) TAT levels were quantified for n=3, and where times of death varied n=5. The data displayed represents means of the duplicate technical repeats plus SDs.

### Delayed drug administration protects against lethality

We initially designed our survival assays in line with the WHO-approved recommendations for the preclinical testing of antivenoms, whereby the preincubation of venom and treatment ensures optimal conditions for the neutralization of venom toxins. To modify this model to better mimic a real-life envenoming scenario, we next challenged groups of mice intraperitoneally with 90 μg (the equivalent of 5 x the intravenous LD_50_ dose) of *E. ocellatus* venom, followed by the administration of an intraperitoneal dose (120 μg) of dimercaprol or DMPS 15 minutes later. All experimental animals were then monitored for signs of envenoming for 24 h, along with those from the venom-only and drug-only control groups. None of the animals receiving metal chelators alone displayed any adverse effects over the 24-h period, while the venom-only group succumbed to the lethal effects of the venom within 1.5-4 h (Fig. 6A). Interestingly, the delayed dosing of DMPS protected against venom-induced lethality for over 12 h after initial envenoming, and two of the five experimental animals survived to the end of the experiment (24 h) (Fig. 6A). The observed prolonged protection against envenoming provided by DMPS is promising, and it is worth noting that the death of the three mice between 12.6-24 h post-venom administration is likely due to the substantial clearance of DMPS in the absence of redosing. Previous research in rats and rabbits has shown that 85-89% of this chelator is excreted in the urine within 6 h following oral dosing *(36, 37)*. Contrastingly, the delayed dosing of dimercaprol in *E. ocellatus*-envenomed mice offered considerably less protection than DMPS. Although two experimental animals also survived until the end of the experiment (24 h), three succumbed rapidly, and at times closely matching those of the venom-only control (111, 125, and 150 min vs. 110, 110, and 143 min, respectively). These findings demonstrate that DMPS outperforms dimercaprol by providing prolonged protection against the lethal effects of *E. ocellatus* snake envenoming in a preclinical model more reminiscent of a real-world envenoming scenario (e.g. treatment delivery after envenoming).

We next investigated whether the delayed administration of DMPS was equally effective against the venoms of *E. pyramidum* and *E. carinatus*. In line with our previous findings, DMPS was more effective at protecting against the lethal effects of *E. pyramidum* venom than those of *E. carinatus*, despite using a higher venom challenge dose (112 vs. 95 μg, respectively) (Fig. 6B and C). While three animals died within the first 6 h (at 99, 227 and 365 min) in the *E. pyramidum* group, two survived for the duration of the experiment (24 h) (Fig. 6B), whereas all mice treated with *E. carinatus* venom succumbed within the first 6.5 h of the experiment, and thus the chelator only provided a slight delay in lethality compared with the venom-only control (Fig. 6C). TAT levels again correlated with lethality, with elevated levels (>400 ng/ml) observed in animals not protected by DMPS, in contrast to those that survived for the duration of the experiment (average of ~200 ng/ml) (Fig. 6D).

**Fig. 6.**
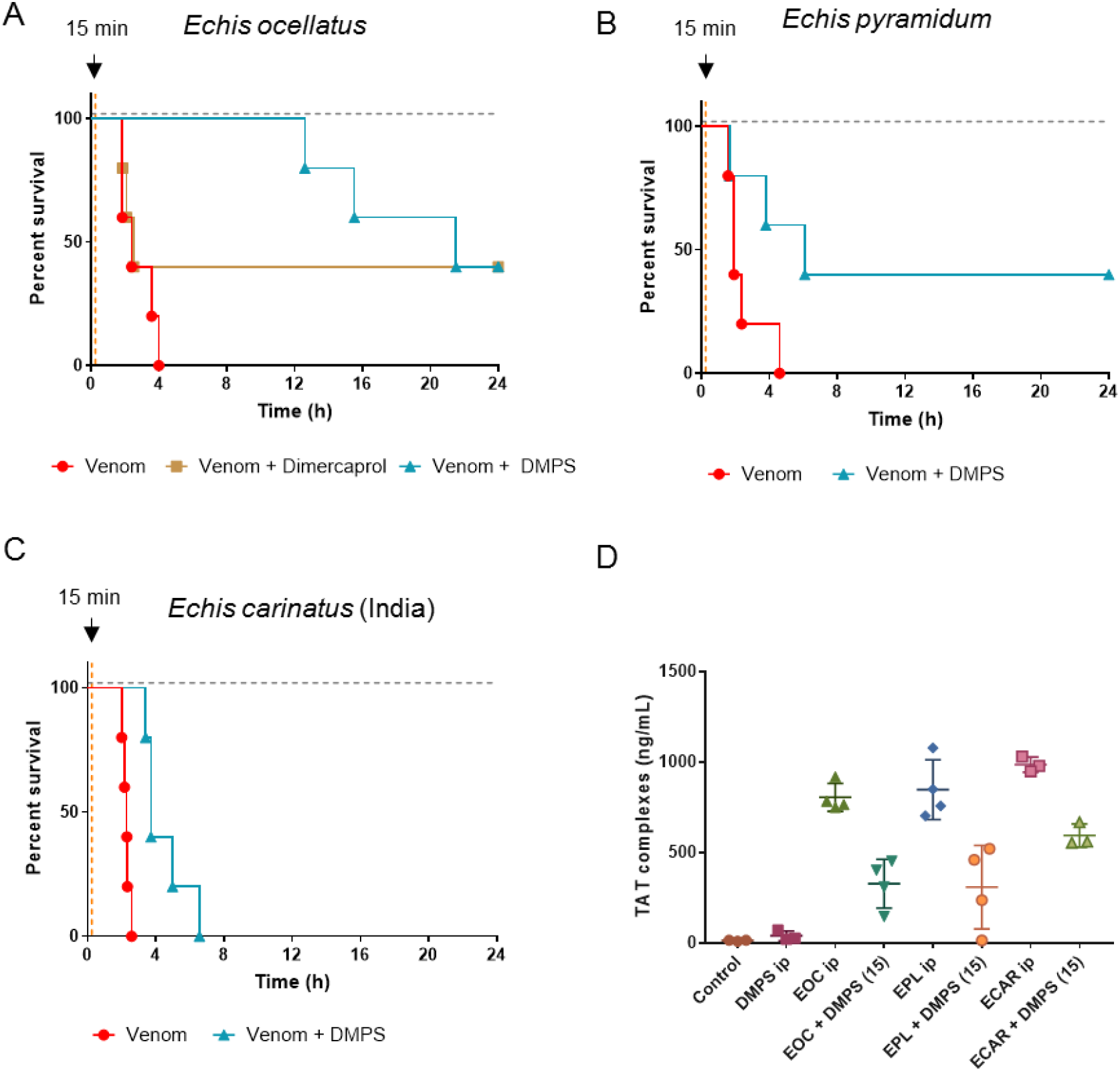
DMPS delays lethality *in vivo* in a ‘challenge and treat’ model of envenoming. Kaplan-Meier survival graphs for experimental animals (n=5) receiving venom (intraperitoneal administration), followed by delayed drug treatment (intraperitoneally 15 min later). (**A**) Survival of mice receiving a 5 x intravenous LD_50_ dose of *E. ocellatus* venom (90 μg) with and without 120 μg of drug (dimercaprol or DMPS) 15 mins later. (**B**) Survival of mice receiving 7 x intravenous LD_50_ dose of *E. pyramidum* venom (112 μg) with and without 120 μg of DMPS 15 mins later. (**C**) Survival of mice receiving 5 x intravenous LD_50_ dose of *E. carinatus* (India) venom (95 μg) with and without 120 μg of DMPS 15 mins later. For (**A**), (**B**) and (**C**) drug-only controls are presented as black dotted lines at the top of each graph (none of the drugs exhibited toxicity at the given doses) and the end of the experiment was at 24 h. (**D**) Quantification of thrombin-antithrombin (TAT) levels in envenomed animals. Where the time of death was the same within experimental groups (e.g. early deaths or complete survival) TAT levels were quantified for n=3, and where times of death varied n=5. The data displayed represents means of the duplicate technical repeats plus SDs.

### DMPS with subsequent antivenom administration prolongs survival

The results described above demonstrate impressive protective capabilities for a single small molecule (DMPS), when considering snake venoms consist of mixtures of numerous (e.g. 50 to >200) toxic constituents. To better advocate for the use of DMPS as an early community-based snakebite therapy, we next sought to model a scenario in which metal chelators were administered as a rapid intervention soon after a snakebite, followed by later antivenom therapy when the patient has reached a healthcare facility. Thus, we compared our earlier findings where *E. ocellatus* venom (90 μg) was administered intraperitoneally followed by DMPS (120 μg) via the same route 15 min later, with mice receiving the same venom and drug regimen, but with the intravenous administration (as per clinical use) of antivenom 1 h post-venom injection (45 mins post-DMPS administration). For this, we used EchiTabG, an ovine antivenom generated against the venom of *E. ocellatus* with known preclinical and clinical efficacy *(13, 38)*. In addition, two control groups were employed: one that received antivenom intravenously 1 h after venom injection but did not receive DMPS, and the other that received antivenom intraperitoneally 15 min after the venom, as a direct comparator for the DMPS dose. A 1-h delay in antivenom administration in the absence of DMPS resulted in the early death of two mice (within the first 4.5 h), which correlated with elevated TAT levels (554 and 728 ng/ml) (Fig. 7A). The antivenom control that was administered intraperitoneally at 15 mins, again in the absence of DMPS, also resulted in one early death (at 142 min) *(39)*. Thus, delayed delivery of high doses of effective antivenom were unable to provide complete protection against envenoming by *E. ocellatus*. Crucially, the early intraperitoneal administration of DMPS at 15 mins post-envenomation, followed by the intravenous delivery of antivenom at 1 h protected against the lethal effects of *E. ocellatus* venom in all experimental animals until the end of the experiment (24 h), thereby extending the survival times of envenomed animals that received DMPS alone by >11 h (first death at 12.6 h) (Fig. 7B). This treatment regimen was also associated with low TAT levels (13.6-23.2 ng/ml), approaching those of the non-envenomed, control mice (10.2-15.9 ng/ml; Fig. 7B). In combination, these findings demonstrate that: (i) DMPS outperforms antivenom therapy when administered by the same route after venom delivery, (ii) DMPS outperforms antivenom therapy when delivered earlier, even when the antivenom is delivered intravenously, and (iii) the combination of early DMPS treatment followed by later antivenom administration provides prolonged protection against *E. ocellatus* venom-induced lethality.

**Fig. 7.**
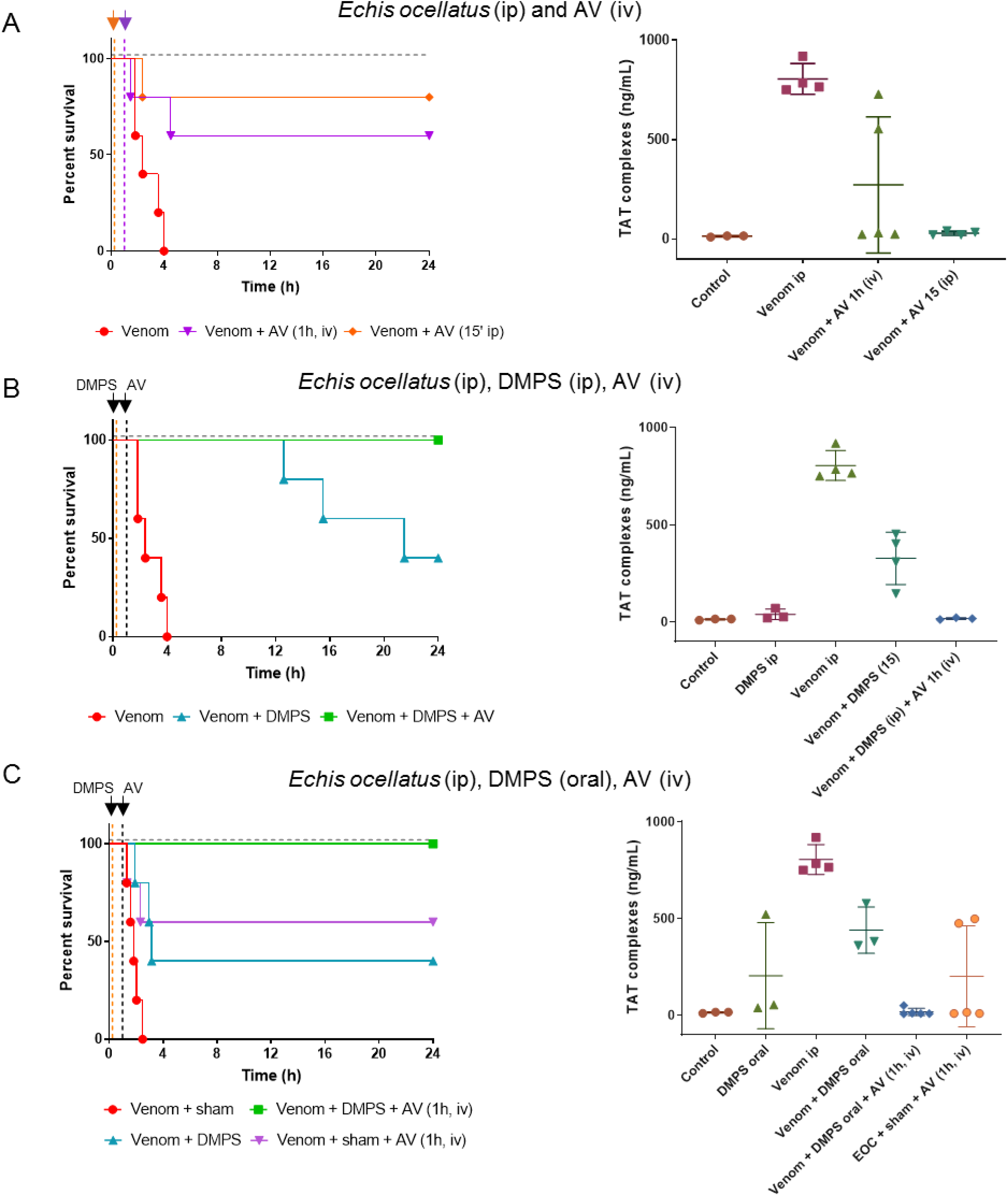
Oral DMPS followed by later administration of antivenom protects against *in vivo* lethality caused by *Echis ocellatus* venom. Kaplan-Meier survival graphs for experimental animals (n=5) receiving *E. ocellatus* venom (90 μg, 5 × intravenous LD_50_ dose, intraperitoneal administration), followed by delayed drug treatment (intraperitoneal or oral) and/or antivenom (intraperitoneal or intravenous). (**A**) Survival of mice receiving *E. ocellatus* venom (intraperitoneally) followed by 168 μl EchiTAbG antivenom either intraperitoneally 15 mins later, or intravenously 1 h later. Corresponding thrombin-antithrombin (TAT) levels for the envenomed animals are depicted on the right. Note: For the venom + AV intraperitoneally (15 min) dataset, the mouse that succumbed early on to the effects of the venom could not be sampled for TAT levels, therefore the data displayed only reflects the animals that survived until the end of the experiment. (**B**) Survival of mice receiving *E. ocellatus* venom (intraperitoneally) followed by DMPS 15 min later (120 μg, intraperitoneally) and antivenom 1 h after venom administration (168 μl, intravenously). Right panel shows corresponding TAT levels. (**C**) Survival of mice receiving *E. ocellatus* venom (intraperitoneally), followed immediately by: (i) oral DMPS (600 μg, ~1 min post-venom injection), (ii) oral DMPS (600 μg, ~1 min post-venom injection) and EchiTAbG antivenom (168 μl, intravenously, 1 hr later), and (iii) EchiTAbG antivenom (168 μl, intravenously, 1 hr later). Right panel shows corresponding TAT levels. For all survival experiments drug- or antivenom-only controls are presented as black dotted lines at the top of each graph (none exhibited toxicity at the given doses) and the end of the experiment was at 24 h. For quantification of TAT, where the time of death was the same within experimental groups (e.g. early deaths or complete survival) TAT levels were quantified for n=3, and where times of death varied n=5. The data displayed represents means of the duplicate technical repeats plus SDs.

### Oral DMPS followed by antivenom protects against venom lethality

DMPS is formulated both as a solution for injection and as an oral capsule, with the latter making it particularly promising as an early intervention for administration outside of a hospital setting. Next, we investigated the preclinical efficacy of DMPS when dosed via the oral route. DMPS was dissolved into a solution containing molasses, which was then provided to experimental animals *ad libitum* via a pipette tip 1 min post-venom injection (90 μg *E. ocellatus* venom administered intraperitoneally; 600 μg DMPS orally; n=5). We compared survival outcomes in experimental groups consisting of: (i) venom and molasses (sham) only, (ii) venom and oral DMPS only, (iii) venom, oral DMPS and then antivenom delivered intravenously 1 h later, and (iv) venom, sham, and the later dose of antivenom (Fig. 7C). The DMPS dose was increased fivefold in these studies (600 μg vs. 120 μg) due to the anticipated lower bioavailability of the oral route compared with the intraperitoneal delivery described earlier. Nonetheless, this dose remains considerably lower than the maximal human equivalent dose permitted over 24 hr (171 mg vs. 2.4 g for a 70 kg adult) or even at a single dosing point (2.44 mg/kg vs 2.86 mg/kg), and the drug itself did not display any signs of toxicity in mice, based on survival data (Fig. 7C) and behavioral and post-mortem observations. Oral DMPS treatment resulted in comparable survival (2 of 5 experimental animals) to intraperitoneally delivered DMPS at the end of the experiment (24 h), although survival times were noticeably prolonged when the drug was delivered via the intraperitoneal route (Figs. 7B and 7C). These findings suggest that, as perhaps anticipated, the uptake and distribution of DMPS via oral administration are indeed inferior to the intraperitoneal route. In the comparator group, where envenomed mice received sham and antivenom only, only three mice survived to the end of the experiment, with two animals succumbing to the lethal effects of the venom (Fig. 7A). Critically though, the combination of oral DMPS followed by later antivenom administration resulted in complete survival in experimental animals at 24 h (Fig. 7C), and those animals displayed TAT levels highly comparable to those observed in normal, non-envenomed, controls (average of 17.3 ng/ml vs. 13.05 ng/ml in the control). These findings demonstrate that the early oral delivery of DMPS is capable of prolonging the survival of envenomed animals, and when combined with a later dose of antivenom, provides prolonged and complete protection against snakebite lethality *in vivo*.

## Discussion

Here, we demonstrate that metal chelators can likely be repurposed as effective snakebite therapeutics, particularly when directed against snake venoms rich in metalloproteinases. Chelators have demonstrated safety profiles, are licensed medicines and show considerable promise as affordable treatments for snakebite in low/middle-income countries (~8 USD/100 mg oral capsule versus 48-315 USD per vial of antivenom *(8)*, with often 5-10 vials required/envenoming). Moreover, the doses at which these SVMP inhibitors show efficacy are >100-fold lower than current immunoglobulin-based treatments (e.g. 60 μg of drug versus 7.5 mg of antivenom for intravenous administration used in this study). The combination of these characteristics make them highly amenable for investigation for clinical use in a community setting, thus potentially dramatically reducing the long time to treatment typically observed after snakebite, which has a known major detrimental impact on patient outcomes *(40, 41)*.

Among the several metal chelators explored in this study, DMPS, a hydrophilic heavy-metal chelator, was effective in preventing lethality *in vivo* in an intravenous model of snakebite envenoming, as well as in prolonging survival in mouse models where the venom challenge was followed by a delayed administration of the drug. Our challenge/treatment model attempted to replicate a clinical situation in which a victim would be bitten, would subsequently receive the metal chelator soon after, and would be later admitted to a healthcare facility where they would be treated with antivenom (if required). In both our models (venom intraperitoneally/DMPS intraperitoneally or venom intraperitoneally/DMPS orally) the delayed administration of DMPS resulted in prolonged animal survival (~17 h and ~10 h compared to the controls for the intraperitoneal and oral dosing, respectively) compared to the venom-only controls. Moreover, when DMPS administration was followed by an intravenous dose of antivenom 1 h later, complete survival at 24 h post-envenoming was observed in all cases, irrespective of whether the drug was delivered intraperitoneally or orally. These findings demonstrate that rapid DMPS intervention substantially delays the onset of venom lethality, as antivenom administration alone (either intraperitoneally as a drug-matched control or intravenously with a 1 h delay) was insufficient to fully neutralize venom activity.

DMPS is approved in Germany for treating mercury intoxication (marketed as Dimaval) and is available as both a 100-mg capsule for oral dosing and a 250-mg ampoule for intravenous or intramuscular administration. In humans, DMPS is readily absorbed following oral administration, can be detected in the plasma 30-45 mins post-administration, and reaches peak plasma concentrations 3-4 h after ingestion *(42)*, making it a strong candidate for an oral community-based therapy. Moreover, the drug forms complexes with plasma proteins via disulfide linkages, which results in slow release and prolongation of its clinical half-life to ~20 h, thus extending its efficacy window *(43)*. Importantly, no major side effects or teratogenic effects have been reported *(44, 45)*, even with repeated oral administration spanning months in animals (126 mg/kg/day in rats for 5 days/week for 66 weeks *(44)*; 45 mg/kg/day over 6 months in Beagles *(46)*) or several days in humans (~15 g DMPS given as a mixed regimen parenterally and orally over 12 days *(47)*). In our study, we used a single therapeutic dose that scales up to 1/14^th^ of the maximal daily oral human equivalent dose regimen for acute heavy metal intoxication (171 mg vs. 2.4 g for a 70 kg adult). However, even under these sub-maximal and single dose conditions, DMPS prolonged survival for at least 12 h in mice envenomed with *E. ocellatus* venom, suggesting that the drug can effectively counteract systemic venom toxicity within a defined therapeutic window. Its efficacy in the intraperitoneal model superseded that of oral administration, likely due to increased bioavailability and distribution, while its decrease in efficacy over time (e.g. >12 h) seems likely to be linked to its excretion in the urine. Previous studies in rats and rabbits indicate that >85% of the drug is eliminated during the first 6 h *(36, 37)*, as opposed to the ~20 h half-life observed in humans. Considering that in our model the drug was only administered once, we hypothesize that repeated dosing may continue to effectively neutralize venom SVMPs, thus expanding the protective interval. We will explore this in future studies.

Of the remaining metal chelators tested in this study, dimercaprol showed some early promise as a snakebite therapeutic. Dimercaprol chelates a variety of heavy metals, including lead, arsenic, gold and mercury, and provided the highest levels of venom inhibition *in vitro* (Figs. 2 and 3). This chelator also neutralized the lethal effects of *E. ocellatus* and *E. carinatus* venoms *in vivo* in the co-incubation model (Figs. 4 and 5), although it showed a lack of efficacy in the more clinically-relevant ‘challenge and treat’ preclinical model we employed (Fig. 6A). In addition, dimercaprol comes with several challenges for clinical use *(48)*, including its formulation in peanut oil, requirement for administration via a deep, painful intramuscular injection, a number of reported adverse reactions, and a small safety margin *(49)*. Given these considerations, its more hydrophilic derivatives offer safer alternatives. Consequently, both DMSA and DMPS have been advocated as the drugs of choice for the contemporary treatment of heavy metal poisoning *(27)*. Our results here, however, demonstrates that DMSA shows a lack of preclinical efficacy against the medically important snake venoms investigated (Fig. 4), thus strongly justifying the selection of DMPS as our lead chelator for future clinical translation.

Our study has several limitations. First, the WHO-recommended protocol for testing snakebite therapies recommends the preincubation of venom with antivenom, which artificially promotes the binding of venom toxins with the therapy. While this is a necessary first step in assessing the therapeutic potential of a treatment by providing a ‘best-case’ scenario, this method does not accurately reflect snakebite, where venom is delivered prior to treatment. To overcome this, we utilized an intraperitoneal challenge/treatment model whereby the treatment was administered after venom, mirroring a more realistic scenario. However, no *in vivo* mouse model, regardless of venom or drug dosage, route of administration, or time between venom injection and drug treatment can ever fully reflect a human snakebite. This is because the venom dose in mice is usually high (45-112 μg) to induce rapid lethality (e.g. <4 h) and avoid prolonged suffering in experimental animals. Contrastingly, lethality from saw-scaled viper bites in humans do not typically occur in <12 h *(30)*. Thus, the onset of pathology is much faster in the animal model, and drug uptake and pharmacokinetics will likely differ due to different circulatory volumes. Nevertheless, the current model is still highly useful in informing antivenom efficacy and is the best preclinical model available for assessing novel therapeutic interventions for snakebite.

Lastly, snake venoms are toxin cocktails *(10)* and their toxin composition can vary extensively among species *(11)*. It is worth noting that DMPS displayed superior efficacy against the venoms of *E. ocellatus* and *E. pyramidum*, compared with the venom of the related species *E. carinatus*, when used as a solo therapy. Nevertheless, DMPS did offer some protection against *E. carinatus* venom, as it was found to prolong survival in experimental animals (Fig. 5B). These differences in efficacy likely reflect variation in the toxin constituents found in the venom of these species (Fig. 1, *(11, 35)*), as a similar lack of preclinical efficacy has been observed when using antivenom made against *E. ocellatus* to neutralize *E. carinatus* venom *(13)* and, conversely, the clinical use of *E. carinatus* antivenom for treating bites by *E. ocellatus* has resulted in poor patient outcomes *(50, 51)*. Despite these limitations, we suggest that DMPS may be a useful early intervention therapeutic to postpone the onset of snakebite pathology caused by a variety of SVMP-rich venoms, particularly when subsequently coupled with species-specific antivenom. We will explore the breadth of DMPS efficacy in combination with antivenom in future work, specifically to assess whether this combination therapy approach may reduce the number of costly vials of antivenom that are currently required to effect cure following snakebite.

Lastly, while we have shown here that DMPS is effective in counteracting the activity of SVMP toxins, it cannot neutralize other toxin classes, such as the enzymatically active serine proteases and PLA_2s_ also found in *E. ocellatus* venom (Fig. S5). Thus, while oral DMPS may still prove to be an effective early intervention for delaying the onset of pathology caused by many viperid snakes, which typically have a high abundance of SVMPs in their venom *(21)*, snakebite victims may still require immediate transport to a healthcare facility in case antivenom therapy is required to neutralize other pathogenic, or even potentially lethal, toxin constituents. Future treatments consisting of mixtures of small molecule inhibitors, each targeting different toxin types, could prove particularly valuable as more generic inhibitors against viper venoms, particularly since such inhibitors with specificities towards different toxin types have recently been described *(15, 52)*.

Despite these described limitations, DMPS remains a strong candidate for translation for clinical use in snakebite envenoming. Given the high tolerance and availability of an oral formulation, DMPS is highly suitable for repurposing for treating snakebite, particularly in the first instance against bites by the West African saw-scaled viper *E. ocellatus*, when considering the preclinical efficacy observed here. This species is an ideal choice for testing the clinical utility of DMPS, as *E. ocellatus* arguably causes more snakebite mortality than any other species of snake. We envision the implementation of a Phase 2 clinical trial in which snakebite victims would be given DMPS as an immediate community-based intervention followed later by antivenom upon arrival at a healthcare facility, compared with a second arm of the trial receiving conventional antivenom therapy only. However, first we plan to undertake a Phase 1 trial to identify an optimal dosing regimen for DMPS, which is necessary because snakebite is an acute event, and thus the optimal bioavailability is likely to differ from that of the current clinical indication of heavy metal poisoning, whereby frequent dosing can be repeated for several weeks. Collectively, our data convincingly demonstrate that inexpensive repurposing of metal chelators can protect against the lethal effects of snakebite. While antivenom may be required secondarily, DMPS seems likely to delay the onset of severe envenoming and may facilitate a reduction in the dose of costly and poorly tolerated antivenom. Ultimately these promising findings strongly advocate for further research into the utility of small molecule inhibitors for rapidly and generically treating the World’s most lethal neglected tropical disease.

## Materials and Methods

### Study design

The study was aimed at determining the efficacy of metal chelators against snakebite *in vitro* and *in vivo*. For all *in vitro* kinetic experiments testing the dose-efficacy of chelators, the assays were performed at least in triplicate and each experiment contained technical triplicates. Within the technical triplicates, outliers were assessed. The exclusion of outliers was performed if one kinetic curve was significantly distinguishable from the other two technical replicates (and considerably shifting the mean) and the mean value of the two ‘good’ replicates was further used in assessing the reproducibility of replicate experiments. For the animal studies, experiments were performed on groups of five, male CD-1 mice, as this is the established WHO protocol for testing the efficacy of anti-snakebite therapies. The mice were randomly distributed in each group and the experimenters assessing the outcomes were blinded to the intervention. Mouse blood was collected form all experimental animals via cardiac puncture, except when this was not possible because the blood had clotted following the intervention. Mouse plasma samples were assessed in duplicate using commercial ELISA kits, with 3<n<5 mice being sampled. The number and sampling for all biological repeats are given in the respective figure legends.

### Statistical analysis

The areas under the curve for kinetic data n=3 were plotted with SEMs. Unpaired two-tailed t-tests were performed in GraphPad Prism 8.1 (GraphPad Software, San Diego, USA) and used to statistically compare the survival times between groups treated with venom alone and venom + drug for both *E. carinatus* and *E. pyramidum*. For all ELISA measurements the biological replicates were plotted with SDs.

### Venoms

Venoms were sourced from either wild-caught specimens maintained in, or historical venom samples stored in, the Herpetarium of the Liverpool School of Tropical Medicine. The venom pools selected encompassed saw-scaled vipers from diverse geographical localities and were from: *E. ocellatus* (Nigeria), *E. carinatus sochureki* (India, referred to throughout as *E. carinatus*), *E. carinatus sochureki* (U.A.E, also referred to as *E. carinatus*), *E. pyramidum leakeyi* (Kenya, referred to as *E. pyramidum*), *E. leucogaster* (Mali) and *E. coloratus* (Egypt). Note that the Indian *E. carinatus* venom was collected from a single specimen that was inadvertently imported to the UK via a boat shipment of stone, and then rehoused at LSTM on the request of the UK RSPCA. Crude venoms were lyophilized and stored at 4 °C to ensure long term stability. Prior to use, venoms were resuspended to 10 mg/ml in PBS (pH 7.4) and then further diluted to 1 mg/ml stock solutions (with PBS) for the described experiments.

### Metal chelators

Dimercaprol (2,3-dimercapto-1-propanol ≥98 % iodometric, Cat no:64046-10 ml), DMSA (meso-2,3-dimercaptosuccinic acid ≥98 % by HPLC, Cat no: D7881-1G) and EDTA (E6758) were purchased from Sigma-Aldrich. DMPS (2,3-dimercapto-1-propanesulfonic acid sodium salt monohydrate, 95%, Cat no: H56578) was sourced from Alfa Aesar. Stocks (tenfold dilutions from 2 mM to 2 μM) were made using deionized water, with the exception of the 2 mM DMSA stock which was made using 100% ethanol, as it is insoluble in water at this concentration. DMPS was resuspended in PBS.

### Venomics

The proteome of *E. leucogaster* (Mali) venom was inferred by comparing the reverse-phase HPLC traces and SDS-PAGE analyses of the chromatographic fractions with those of the previously reported *(11)* venom proteomes of *E. ocellatus* (Nigeria), *E. carinatus* (U.A.E. and India), *E. pyramidum* (Kenya), and *E. coloratus* (Egypt), and the partially characterized venom proteome of *E. leucogaster* (Mali) *(53)*. The relative abundances (expressed as percentage of the total venom proteins) of the different protein families were calculated from the relation of the sum of the areas of the reverse-phase chromatographic peaks (containing proteins from the same toxin family), to the total area of venom protein peaks in the reverse-phase chromatogram. The relative contributions of different proteins eluting in the same chromatographic fraction was estimated by densitometry of Coomassie brilliant blue-stained SDS-PAGE gels, as previously outlined *(54)*.

### SVMP assay

The SVMP activity of the various venoms, in the presence or absence of inhibitors, was measured using a quenched fluorogenic substrate (ES010, R&D Biosystems). Briefly, 10 μl of the substrate (supplied as a 6.2 mM stock) was used per 5 ml reaction buffer (150 mM NaCl, 50 mM Tris-Cl pH 7.5). Reactions consisted of 10 μl of venom ± inhibitors in PBS and 90 μl of substrate. Venoms were used at 1 μg/reaction and the final concentrations of the various inhibitors ranged from 150 μM to 150 nM (tenfold dilutions). The venom and inhibitors were preincubated for 30 min at 37 °C and pipetted in triplicate onto 384-well plates (Greiner). A Labsystems Multidrop Reagent dispenser was used to dispense the substrate. The plate was run on an Omega FluoSTAR (BMG Labtech) instrument at an excitation wavelength of 320 nm and emission wavelength of 405 nm at 25 °C for 1 h. The areas under the curve (AUCs) in the 0-40 min interval were calculated for each sample; this time point was chosen as the time where all fluorescence curves had typically reached a plateau (maximum fluorescence). For comparing venom-only samples, the averages of at least three independent experimental runs for each condition, expressed as AUCs (n ≥ 3), were plotted at each inhibitor concentration with standard error of the mean (S.E.M). To determine inhibitor efficacy, the AUCs for each of the samples that consisted of venom + inhibitors were transformed and expressed as percentages of the venom-only sample (where the venom was 100%). The negative control (PBS only) was also expressed relative to the venom and the variation in background levels was presented as an interval (dotted green lines) delineated by the lowest and highest value in the PBS-only sample across concentrations and inhibitors for a specific venom.

### Plasma assay

To assess the pro- or anti-coagulant activity of our venoms, we used a previously published method *(33)*. Briefly, 100 ng of each venom was incubated at 37 °C for 30 min in the presence or absence of inhibitors, whose concentrations ranged from 150 μM to 150 nM (tenfold dilutions). The final reaction consisted of 10 μl venom ± inhibitors in PBS, 20 μl 20 mM CaCl2, and 20 μl citrated bovine plasma (VWR). The samples were pipetted in triplicate onto 384-well plates and the absorbance was monitored at 595 nm for 2 h at 25 °C on an Omega FluoSTAR instrument (kinetic cycle ~68 s). The time point where the plasma control intersected the venom-only sample (the time required for the plasma to fully clot, Fig. S6) was determined in each case. We next generated the AUCs for the 0-n time interval, where n represents the number of minutes required for the plasma control to clot. Of note, we have observed that differences in the plasma clotting time influenced by the presence of venoms can affect the ‘height’ of the plateau - i.e. the final absorbance at which a plasma sample clots was usually higher than that of the venom-only sample. As the presence of the venom in a sample will lead to faster coagulation, we presume this may influence the formation of the clot, resulting in clots through which light passes differently. To avoid artificial increases in the calculated AUCs due to differences in plateau height, we transformed the data so that the venom-only maximal absorbance was applied to all venom + inhibitor samples for the respective dataset (see Fig. S6). Therefore, any shifts in clotting (e.g. increased inhibition associated with a shift to the right closer to the plasma control; less inhibition associated with a shift to the left, closer to the venom-only control) would be due only to the ability of the inhibitor to counteract the procoagulant nature of the venom. To compare total venom activity, the averages of at least three independent experimental runs for each condition, expressed as AUCs (n ≥ 3), were plotted at each inhibitor concentration with S.E.Ms. To determine inhibitor efficacy, the AUCs for each of the samples that consisted of venom + inhibitors were transformed and expressed as percentages of the venom-only sample (where the venom was 100%). The negative control (plasma only) was also expressed relative to the venom and the variation in background levels were presented as an interval (dotted black lines) delineated by the lowest and highest value in the plasma-only sample across concentrations and inhibitors for a specific venom.

### Prothrombin degradation

To assess the ability of the venoms to degrade prothrombin, we incubated 5 μg of venom with 5 μg of prothrombin (Haematological Technologies, Inc) in the presence or absence of inhibitors (at 150 and 500 μM) in a final volume of 15 μl. Venoms and inhibitors were preincubated at 37 °C for 30 min, after which prothrombin was added, followed by another 10 min incubation at 37 °C. The reaction was stopped by adding an equivalent volume of 2X PAGE-loading dye containing β-mercaptoethanol, after which the samples were heated at 100 °C for 5 min. The entire sample volume (30 μl) was loaded and run on 4-20% 12-well Novex precast gels (Thermo Fisher) and visualized with Coomassie Brilliant Blue.

### Preclinical studies

All animal experiments were conducted using protocols approved by the Animal Welfare and Ethical Review Boards of the Liverpool School of Tropical Medicine and the University of Liverpool, and performed in specific pathogen-free conditions under licensed approval of the UK Home Office and in accordance with the Animal [Scientific Procedures] Act 1986 and institutional guidance on animal care. Experimental design was based upon refined WHO-recommended protocols *(8, 55)*, with the observers being blinded to the experimental groups.

The median lethal dose (venom LD_50_) for *E. ocellatus* (Nigeria), *E. carinatus* (India) and *E. pyramidum* (Kenya) venoms (18, 19, and 16 μg/20 g mouse respectively) were previously determined *(8, 24, 55)*. Drug stocks were freshly prepared as follows: DMPS (1 mg/ml in PBS), DMSA (2 mg/ml in ethanol, with a final concentration of 15% for the experimental dose) and Dimercaprol (1 mg/ml in water).

### Co-incubation model of preclinical efficacy

For our initial experiments, we used 2.5 x the intravenous LD_50_ doses of *E. ocellatus* (45 μg), *E. carinatus* (India) (47.5 μg) and *E. pyramidum* venoms (40 μg) in a refined version of the WHO recommended *(55)* antivenom ED_50_ neutralization experiments *(24)*. Groups of five male 18-22 g CD-1 mice (Charles River, UK) received experimental doses that consisted of either (a) venom only (2.5 x LD_50_ dose); (b) venom and drug (60 μg); or (c) drug only (60 μg). All experimental doses were prepared to a volume of 200 μl in PBS and incubated at 37 °C for 30 mins prior to their intravenous injection via the tail vein. Animals were monitored for 6 h and euthanized upon observation of humane endpoints (seizure, pulmonary distress, paralysis, hemorrhage). Deaths, time of death, and survivors were recorded; where “deaths/time of death” actually represents the implementation of euthanasia based on the defined humane endpoints.

### Challenge and treatment model of preclinical efficacy

Next, we performed experiments whereby mice were initially challenged with venom, followed by delayed administration of the drug/antivenom. For these studies, we used an intraperitoneal model of venom administration to provide an acceptable time window to measure venom neutralization. Consequently, the venom challenge doses were increased to 5 x the intravenous LD_50_s (*E. ocellatus* [90 μg] and *E. carinatus* India [95 μg]) or 7 intravenous LD_50_s (*E. pyramidum leakeyi* [112 μg]) to ensure complete lethality in the venom-only control group occurred within 4-5 h. Groups of five male 18-22 g CD-1 mice (Charles River, UK) were injected with venom (100 μl final volume), followed by a drug dose that was scaled up accordingly (120 μg + PBS up to 200 μl) after 15 min. The experimental groups comprised mice receiving: (a) venom only (5 x intravenous LD_50_ or 7 x intravenous LD_50_) + 200 μl PBS (15 min later); (b) venom (5 x intravenous LD_50_ or 7 x intravenous LD_50_) + drug (120 μg, 15 min later); and (c) sham (100 μl PBS) + drug (120 μg, 15 min later).

For later ‘challenge and treat’ experiments with *E. ocellatus* venom only, we also explored the efficacy of antivenom, both with and without drug treatment. The antivenom used was EchiTAbG (MicroPharm Limited, UK), an ovine monospecific anti-E. *ocellatus* antivenom validated for preclinical and clinical efficacy against envenomings by this species *(13, 38)*. The median effective dose (ED_50_) of EchiTAbG (Micropharm, U.K.) against 5 x LD_50_ *E. ocellatus* (intravenous LD_50_ of 12.43 μg/mouse) venom was previously determined to be 58.46 μl *(13)*. Consequently, we scaled up the antivenom dose to a dose that would protect all envenomed animals (2 x ED_50_; 168 μl), taking into account the slight difference in LD_50_ of our current *E. ocellatus* venom batch (17.85 μg/mouse). Antivenom was administered either intravenously 1 h post-venom injection or intraperitoneally as a drug-matched control after 15 mins. The experimental design followed that described above with the following experimental groups: (a) *E. ocellatus* venom (90 μg; 5 x intravenous LD_50_) + drug (120 μg, 15 min later) + antivenom (intravenously 1 h later, 168 μl); (b) *E. ocellatus* venom (90 μg; 5 x LD_50_) + antivenom (intravenously, 1 h later, 168 μl); (c) *E. ocellatus* venom (5 x intravenous LD_50_) + antivenom (intraperitoneally 15 min later, 168 μl). For both sets of experiments described above, experimental animals were monitored for 24 h and euthanized upon observation of humane endpoints (seizure, pulmonary distress, paralysis, hemorrhage). Deaths, time of death, and survivors were recorded; where “deaths/time of death” represents the implementation of euthanasia based on the defined humane endpoints. Unpaired two-tailed t-tests in GraphPad Prism 8.1 (GraphPad Software, San Diego, USA) were used to compare the survival times between groups treated with venom alone and venom + drug for *E. carinatus* and *E. pyramidum* venoms.

### Challenge and treatment model using oral DMPS

Next, we assessed the utility of DMPS as an oral therapeutic, using a similar ‘challenge and treat’ approach. We injected groups of mice intraperitoneally with 5 x intravenous LD_50_s of *E. ocellatus* venom, followed immediately (~1 min) by an oral dose of DMPS (600 μg, a dose lower than the human equivalent dose *(56)* [e.g. 171 mg vs. 2.4 g maximal daily dose permitted for a 70 kg adult, or 2.44 mg/kg vs 2.86 mg/kg every 2 h]). The drug was dissolved in 50 μl of ~50 mg/ml molasses and administered orally (*ad libitum*) via a pipette tip to groups of 5 male CD-1 mice. The experimental groups comprised: (a) venom (90 μg) + molasses; (b) venom (90 μg) + oral DMPS (600 μg); (c) venom (90 μg) + oral DMPS (600 μg) + antivenom (intravenously, 1 h later, 168 μl); (d) venom (90 μg) + molasses + antivenom (intravenously, 1 h later, 168 μl). Animals were monitored for 24 h and euthanized upon observation of humane endpoints, as previously described.

### Thrombin-antithrombin ELISA

For all experimental animals described above, blood samples were collected via cardiac puncture immediately post-euthanasia. Plasma was separated by centrifugation at 400 x *g* for 10 min and stored at −80 °C. We assessed the levels of thrombin-antithrombin complexes (TAT) using a mouse ELISA Kit (Abcam, ab137994), following the manufacturer’s protocol. Mouse plasma samples were diluted 1:100 or 1:150 to fit within the linear range of the assay and measurements were performed in duplicate. All available plasma samples (some were unobtainable via cardiac puncture due to extensive internal hemorrhage) were assessed if the time of death within the group varied, whereas three samples were randomly chosen if the time of death was the same (e.g. either very rapid death within 2 minutes, or survival until the end of the experiment ~360 min or 24 h). The resulting data was plotted as the median of duplicate measurements for each animal and is presented with standard deviations (SDs).

### Echis ocellatus venom fractionation

Proteins from *E. ocellatus* venom (50 mg in 5 ml of PBS) were separated using gel filtration chromatography on a Superdex 200 matrix packed in a 2.6 x 100 cm (500 ml) column. The column was run at 2 ml/min in PBS and elution was monitored at 280 nm. Fractions of 6 ml were collected and the protein content analyzed by SDS-PAGE under reducing conditions using 4-20% gradient gels as above [see Prothrombin Degradation]. Key fractions were assayed for serine protease and PLA_2_ activity, as described below.

### Serine protease assay

To determine the serine protease activity of the various *E. ocellatus* venom fractions, we used a chromogenic kinetic assay, and the specific substrate S-2288 (Cambridge Biosciences). Samples (5 μl of each fraction) were plated onto 384-well plates, and then overlaid with buffer (100 mM Tris-Cl, 100 mM NaCl, pH 8.5), and 6 mM of S-2288. Changes in absorbance were measured at 405 nm for ~30 mins. Negative control readings (PBS) were subtracted from each reading and the rate of substrate consumption calculated by measuring the slope between 0 and 5 mins. To assess inhibition by DMPS, samples that displayed activity were preincubated with DMPS at a final concentration of 150 μM for 30 min at 37 °C and assayed as above. The means of duplicate or triplicate measurements with SDs were plotted.

### PLA_2_ assay

To assess PLA_2_ activity, we used the EnzChek™ Phospholipase A2 Assay Kit (#E10217, Fisher Scientific), following the manufacturer’s instructions. Briefly, 5 μl of the various venom fractions were assayed and compared with the bee venom standard, alongside a negative control containing no venom. For testing DMPS efficacy in the fractions which displayed PLA_2_ activity, DMPS at a final concentration of 150 μM was preincubated with the fractions for 30 min at 37 °C and a drug-only control was also measured. A standard activity curve was generated using 5, 4, 3, 2, 1 and 0 U/ml of bee PLA_2_ enzyme present in the kit. Fifty microliter samples were mixed with 50 μl of substrate mix and the reaction was incubated in the dark for 10 mins. End-point fluorescence was then measured on a FLUOStar Omega Instrument at an excitation wavelength of 485 nm and an emission wavelength of 520 nm. The negative control was subtracted from the raw values for each sample and PLA_2_ activity was calculated (U/ml) in each fraction. The means of duplicate or triplicate experimental measurements with SDs were plotted.

## Acknowledgments

We thank Paul Rowley for maintenance and husbandry of the snake collection at LSTM and for performing venom extractions.

## Author contributions

L-O.A., J.K. and N.R.C designed the study. L-O.A. and M.H. performed *in vitro* tests. J.J.C. performed proteomics on *E. leucogaster* venom. M.W. performed fractionation of *E. ocellatus* venom. S.A., J.A., E.C., R.A.H. and N.R.C. performed the *in vivo* research. L-O.A. wrote the manuscript with input from J.K. and N.R.C.

## Supplementary Materials

Fig. S1. The snake venom metalloproteinase and plasma clotting activity of saw-scaled viper venoms.

Fig. S2. Metal chelators inhibit the procoagulant activity of saw-scaled viper venoms in the absence of calcium.

Fig. S3. Degradation of prothrombin by saw-scaled viper venoms and inhibition of this activity by metal chelators.

Fig. S4. Correlations between in vivo survival and thrombin-antithrombin levels.

Fig. S5. DMPS does not inhibit serine protease and PLA_2_ activities in *E. ocellatus* venom.

Fig. S6. Processing of plasma clotting data.

## References

1. J. M. Gutiérrez, J. J. Calvete, A. G. Habib, R. A. Harrison, D. J. Williams, D. A. Warrell, Snakebite envenoming, Nat. Rev. Dis. Prim. 3, 17063 (2017).

2. A. G. Habib, A. Kuznik, M. Hamza, M. I. Abdullahi, B. A. Chedi, J.-P. Chippaux, D. A. Warrell, H. J. de Silva, Ed. Snakebite is Under Appreciated: Appraisal of Burden from West Africa, PLoS Negl. Trop. Dis. 9, e0004088 (2015).

3. R. A. Harrison, A. Hargreaves, S. C. Wagstaff, B. Faragher, D. G. Lalloo, Snake envenoming: A disease of poverty, PLoS Negl. Trop. Dis. 3 (2009), doi:10.1371/journal.pntd.0000569.

4. J. Longbottom, F. M. Shearer, M. Devine, G. Alcoba, F. Chappuis, D. J. Weiss, S. E. Ray, N. Ray, D. A. Warrell, R. Ruiz de Castañeda, D. J. Williams, S. I. Hay, D. M. Pigott, Vulnerability to snakebite envenoming: a global mapping of hotspots., Lancet (London, England) 392, 673–684 (2018).

5. B. Mohapatra, D. A. Warrell, W. Suraweera, P. Bhatia, N. Dhingra, R. M. Jotkar, P. S. Rodriguez, K. Mishra, R. Whitaker, P. Jha, for the M. D. S. Collaborators, J. O. Gyapong, Ed. Snakebite Mortality in India: A Nationally Representative Mortality Survey, PLoS Negl. Trop. Dis. 5, e1018 (2011).

6. D. J. Williams, J.-M. Gutiérrez, J. J. Calvete, W. Wüster, K. Ratanabanangkoon, O. Paiva, N. I. Brown, N. R. Casewell, R. A. Harrison, P. D. Rowley, M. O’Shea, S. D. Jensen, K. D. Winkel, D. A. Warrell, Ending the drought: New strategies for improving the flow of affordable, effective antivenoms in Asia and Africa, J. Proteomics 74, 1735–1767 (2011).

7. H. Khosrojerdi, M. Amini, Acute and Delayed Stress Symptoms Following Snakebite, Mashhad Univ. Med. Sci. 2, 140–144 (2013).

8. R. A. Harrison, G. O. Oluoch, S. Ainsworth, J. Alsolaiss, F. Bolton, A.-S. Arias, J.-M. Gutiérrez, P. Rowley, S. Kalya, H. Ozwara, N. R. Casewell, J.-P. Chippaux, Ed. Preclinical antivenom-efficacy testing reveals potentially disturbing deficiencies of snakebite treatment capability in East Africa, PLoS Negl. Trop. Dis. 11, e0005969 (2017).

9. D. J. Williams, M. A. Faiz, B. Abela-Ridder, S. Ainsworth, T. C. Bulfone, A. D. Nickerson, A. G. Habib, T. Junghanss, H. W. Fan, M. Turner, R. A. Harrison, D. A. Warrell, J. M. Gutiérrez, Ed. Strategy for a globally coordinated response to a priority neglected tropical disease: Snakebite envenoming, PLoS Negl. Trop. Dis. 13, e0007059 (2019).

10. N. R. Casewell, W. Wüster, F. J. Vonk, R. A. Harrison, B. G. Fry, Complex cocktails: the evolutionary novelty of venoms, Trends Ecol. Evol. 28, 219–229 (2013).

11. N. R. Casewell, S. C. Wagstaff, W. Wuster, D. A. N. Cook, F. M. S. Bolton, S. I. King, D. Pla, L. Sanz, J. J. Calvete, R. A. Harrison, Medically important differences in snake venom composition are dictated by distinct postgenomic mechanisms, Proc. Natl. Acad. Sci. 111, 9205–9210 (2014).

12. J. P. Chippaux, V. Williams, J. White, Snake venom variability: methods of study, results and interpretation., Toxicon 29, 1279–303 (1991).

13. N. R. Casewell, D. A. N. Cook, S. C. Wagstaff, A. Nasidi, N. Durfa, W. Wüster, R. A. Harrison, D. J. Williams, Ed. Pre-Clinical Assays Predict Pan-African Echis Viper Efficacy for a Species-Specific Antivenom, PLoS Negl. Trop. Dis. 4, e851 (2010).

14. H. A. de Silva, A. Pathmeswaran, C. D. Ranasinha, S. Jayamanne, S. B. Samarakoon, A. Hittharage, R. Kalupahana, G. A. Ratnatilaka, W. Uluwatthage, J. K. Aronson, J. M. Armitage, D. G. Lalloo, H. J. de Silva, K. Winkel, Ed. Low-Dose Adrenaline, Promethazine, and Hydrocortisone in the Prevention of Acute Adverse Reactions to Antivenom following Snakebite: A Randomised, Double-Blind, Placebo-Controlled Trial, PLoS Med. 8, e1000435 (2011).

15. T. C. Bulfone, S. P. Samuel, P. E. Bickler, M. R. Lewin, Developing Small Molecule Therapeutics for the Initial and Adjunctive Treatment of Snakebite, J. Trop. Med. 2018, 1–14 (2018).

16. M. Lewin, L. Gilliam, J. Gilliam, S. Samuel, T. Bulfone, P. Bickler, J. Gutiérrez, Delayed LY333013 (Oral) and LY315920 (Intravenous) Reverse Severe Neurotoxicity and Rescue Juvenile Pigs from Lethal Doses of Micrurus fulvius (Eastern Coral Snake) Venom, Toxins (Basel). 10, 479 (2018).

17. M. Lewin, J. Gutiérrez, S. Samuel, M. Herrera, W. Bryan-Quirós, B. Lomonte, P. Bickler, T. Bulfone, D. Williams, Delayed Oral LY333013 Rescues Mice from Highly Neurotoxic, Lethal Doses of Papuan Taipan (Oxyuranus scutellatus) Venom, Toxins (Basel). 10, 380 (2018).

18. Y. Wang, J. Zhang, D. Zhang, H. Xiao, S. Xiong, C. Huang, Exploration of the Inhibitory Potential of Varespladib for Snakebite Envenomation, Molecules 23, 391 (2018).

19. S. Takeda, H. Takeya, S. Iwanaga, Snake venom metalloproteinases: Structure, function and relevance to the mammalian ADAM/ADAMTS family proteins, Biochim. Biophys. Acta - Proteins Proteomics 1824, 164–176 (2012).

20. R. Kini, C. Koh, R. M. Kini, C. Y. Koh, Metalloproteases Affecting Blood Coagulation, Fibrinolysis and Platelet Aggregation from Snake Venoms: Definition and Nomenclature of Interaction Sites, Toxins (Basel). 8, 284 (2016).

21. T. Tasoulis, G. K. Isbister, A Review and Database of Snake Venom Proteomes., Toxins (Basel). 9, 290 (2017).

22. A. S. Arias, A. Rucavado, J. M. Gutiérrez, Peptidomimetic hydroxamate metalloproteinase inhibitors abrogate local and systemic toxicity induced by Echis ocellatus (saw-scaled) snake venom, Toxicon 132, 40–49 (2017).

23. A. Rucavado, T. Escalante, A. Franceschi, F. Chaves, G. León, Y. Cury, M. Ovadia, J. M. Gutiérrez, Inhibition of local hemorrhage and dermonecrosis induced by Bothrops asper snake venom: effectiveness of early in situ administration of the peptidomimetic metalloproteinase inhibitor batimastat and the chelating agent CaNa2EDTA., Am. J. Trop. Med. Hyg. 63, 313–319 (2000).

24. S. Ainsworth, J. Slagboom, N. Alomran, D. Pla, Y. Alhamdi, S. I. King, F. M. S. Bolton, J. M. Gutiérrez, F. J. Vonk, C.-H. Toh, J. J. Calvete, J. Kool, R. A. Harrison, N. R. Casewell, The paraspecific neutralisation of snake venom induced coagulopathy by antivenoms, Commun. Biol. 1, 34 (2018).

25. J. M. Howes, R. D. G. Theakston, G. D. Laing, Neutralization of the haemorrhagic activities of viperine snake venoms and venom metalloproteinases using synthetic peptide inhibitors and chelators, Toxicon 49, 734–739 (2007).

26. A. N. Nanjaraj Urs, M. Yariswamy, C. Ramakrishnan, V. Joshi, K. N. Suvilesh, M. N. Savitha, D. Velmurugan, B. S. Vishwanath, Inhibitory potential of three zinc chelating agents against the proteolytic, hemorrhagic, and myotoxic activities of Echis carinatus venom, Toxicon 93, 68–78 (2015).

27. S. J. S. Flora, V. Pachauri, Chelation in metal intoxication, Int. J. Environ. Res. Public Health 7, 2745–2788 (2010).

28. D. A. Warrell, C. Arnett, The importance of bites by the saw-scaled or carpet viper (Echis carinatus): epidemiological studies in Nigeria and a review of the world literature., Acta Trop. 33, 307–41 (1976).

29. R. N.. Pugh, R. D.. Theakston, Incidence and mortality of snakebite in savanna Nigeria, Lancet 316, 1181–1183 (1980).

30. D. A. Warrell, Davidson NMcD, B. M. Greenwood, L. D. Ormerod, H. M. Pope, B. J. Watkins, C. R. Prentice, Poisoning by bites of the saw-scaled or carpet viper (Echis carinatus) in Nigeria., Q. J. Med. 46, 33–62 (1977).

31. D. Bartholdi, C. Selic, J. Meier, H. Jung, Viper snakebite causing symptomatic intracerebral haemorrhage, J. Neurol. 251, 889–891 (2004).

32. M. J. Kosnett, The Role of Chelation in the Treatment of Arsenic and Mercury Poisoning, J. Med. Toxicol. 9, 347–354 (2013).

33. K. Still, R. Nandlal, J. Slagboom, G. Somsen, N. Casewell, J. Kool, K. B. M. Still, R. S. S. Nandlal, J. Slagboom, G. W. Somsen, N. R. Casewell, J. Kool, Multipurpose HTS Coagulation Analysis: Assay Development and Assessment of Coagulopathic Snake Venoms, Toxins (Basel). 9, 382 (2017).

34. A. Rogalski, C. Soerensen, B. Op den Brouw, C. Lister, D. Dashevsky, K. Arbuckle, A. Gloria, C. N. Zdenek, N. R. Casewell, J. M. Gutiérrez, W. Wüster, S. A. Ali, P. Masci, P. Rowley, N. Frank, B. G. Fry, Differential procoagulant effects of saw-scaled viper (Serpentes: Viperidae: Echis) snake venoms on human plasma and the narrow taxonomic ranges of antivenom efficacies., Toxicol. Lett. 280, 159–170 (2017).

35. N. R. Casewell, R. A. Harrison, W. Wüster, S. C. Wagstaff, Comparative venom gland transcriptome surveys of the saw-scaled vipers (Viperidae: Echis) reveal substantial intra-family gene diversity and novel venom transcripts, BMC Genomics 10, 564 (2009).

36. B. Gabard, Distribution and excretion of the mercury chelating agent sodium 2,3-dimercaptopropane-1-sulfonate in the rat, Arch. Toxicol. 39, 289–298 (1978).

37. N. I. Luganskii, I. I. Loboda, [Conversion of unithiol in the organism]., Farmakol. Toksikol. 23, 349–55 (1960).

38. I. S. Abubakar, S. B. Abubakar, A. G. Habib, A. Nasidi, N. Durfa, P. O. Yusuf, S. Larnyang, J. Garnvwa, E. Sokomba, L. Salako, R. D. G. Theakston, E. Juszczak, N. Alder, D. A. Warrell, for the N.-U. E. S. Group, D. G. Lalloo, Ed. Randomised Controlled Double-Blind Non-Inferiority Trial of Two Antivenoms for Saw-Scaled or Carpet Viper (Echis ocellatus) Envenoming in Nigeria, PLoS Negl. Trop. Dis. 4, e767 (2010).

39. Y. Mise, R. Lira-da-Silva, F. Carvalho, Time to treatment and severity of snake envenoming in Brazil, Rev. Panam. Salud Pública 42, 1–6 (2018).

40. S. B. Abubakar, A. G. Habib, J. Mathew, Amputation and disability following snakebite in Nigeria, Trop. Doct. 40, 114–116 (2010).

41. S. K. Sharma, F. Chappuis, N. Jha, P. A. Bovier, L. Loutan, S. Koirala, Impact of snake bites and determinants of fatal outcomes in southeastern Nepal., Am. J. Trop. Med. Hyg. 71, 234–8 (2004).

42. R. M. Maiorino, R. C. Dart, D. E. Carter, H. V Aposhian, Determination and metabolism of dithiol chelating agents. XII. Metabolism and pharmacokinetics of sodium 2,3-dimercaptopropane-1-sulfonate in humans., J. Pharmacol. Exp. Ther. 259, 808–14 (1991).

43. R. M. Maiorino, Z. F. Xu, H. V Aposhian, Determination and metabolism of dithiol chelating agents. XVII. In humans, sodium 2,3-dimercapto-1-propanesulfonate is bound to plasma albumin via mixed disulfide formation and is found in the urine as cyclic polymeric disulfides., J. Pharmacol. Exp. Ther. 277, 375–84 (1996).

44. F. Planas-Bohne, B. Gabard, E. H. Schäffer, Toxicological studies on sodium 2,3-dimercaptopropane-1-sulfonate in the rat., Arzneimittelforschung. 30, 1291–4 (1980).

45. J. L. Domingo, A. Ortega, M. A. Bosque, J. Corbella, Evaluation of the developmental effects on mice after prenatal, or pre- and postnatal exposure to 2,3-dimercaptopropane-1-sulfonic acid (DMPS)., Life Sci. 46, 1287–92 (1990).

46. L. Szinicz, P. Wiedemann, H. Häring, N. Weger, Effects of repeated treatment with sodium 2,3-dimercaptopropane-1-sulfonate in beagle dogs., Arzneimittelforschung. 33, 818–21 (1983).

47. R. Heinrich-Ramm, K. Schaller, J. Horn, J. Angerer, Arsenic species excretion after dimercaptopropanesulfonic acid (DMPS) treatment of an acute arsenic trioxide poisoning, Arch. Toxicol. 77, 63–68 (2003).

48. J. Aaseth, M. A. Skaug, Y. Cao, O. Andersen, Chelation in metal intoxication-Principles and paradigms, J. Trace Elem. Med. Biol. 31, 260–266 (2015).

49. American Society of Hospital Pharmacists., AHFS drug information 2010. (American Society of Health-System Pharmacists, Bethesda, MD:, 2010; https://www.worldcat.org/title/ahfs-drug-information-2010/oclc/609401493).

50. E. Alirol, P. Lechevalier, F. Zamatto, F. Chappuis, G. Alcoba, J. Potet, H. J. de Silva, Ed. Antivenoms for Snakebite Envenoming: What Is in the Research Pipeline?, PLoS Negl. Trop. Dis. 9, e0003896 (2015).

51. L. E. Visser, S. Kyei-Faried, D. W. Belcher, D. W. Geelhoed, J. S. van Leeuwen, J. van Roosmalen, Failure of a new antivenom to treat Echis ocellatus snake bite in rural Ghana: the importance of quality surveillance, Trans. R. Soc. Trop. Med. Hyg. 102, 445–450 (2008).

52. L.-O. Albulescu, T. Kazandjian, J. Slagboom, B. Bruyneel, S. Ainsworth, J. AlSolaiss, S. C. Wagstaff, G. Whiteley, R. A. Harrison, C. Ulens, J. Kool, N. R. Casewell, A decoy-receptor approach using nicotinic acetylcholine receptor mimics reveals their potential as novel therapeutics against neurotoxic snakebite, Front. Pharmacol. 10, 848 (2019).

53. J. J. Calvete, P. Cid, L. Sanz, Á. Segura, M. Villalta, M. Herrera, G. León, R. Harrison, N. Durfa, A. Nasidi, R. D. G. Theakston, D. A. Warrell, J. M. Gutiérrez, Antivenomic Assessment of the Immunological Reactivity of EchiTAb-Plus-ICP, an Antivenom for the Treatment of Snakebite Envenoming in Sub-Saharan Africa, Am. J. Trop. Med. Hyg. 82, 1194–1201 (2010).

54. S. Eichberg, L. Sanz, J. J. Calvete, D. Pla, Constructing comprehensive venom proteome reference maps for integrative venomics, Expert Rev. Proteomics 12, 557–573 (2015).

55. R. D. Theakston, H. A. Reid, Development of simple standard assay procedures for the characterization of snake venom., Bull. World Health Organ. 61, 949–56 (1983).

56. A. B. Nair, S. Jacob, A simple practice guide for dose conversion between animals and human, J. Basic. Clin. Pharm. 7(2), 27–31.

57. A. Patra, B. Kalita, A. Chanda, A. K. Mukherjee, Proteomics and antivenomics of Echis carinatus carinatus venom: Correlation with pharmacological properties and pathophysiology of envenomation, Sci. Rep. 7, 1–17 (2017).

