## Supplemental Figures 1-6 for "Preclinical validation of a repurposed metal chelator as a community-based therapeutic for hemotoxic snakebite"

**Supplementary Materials**

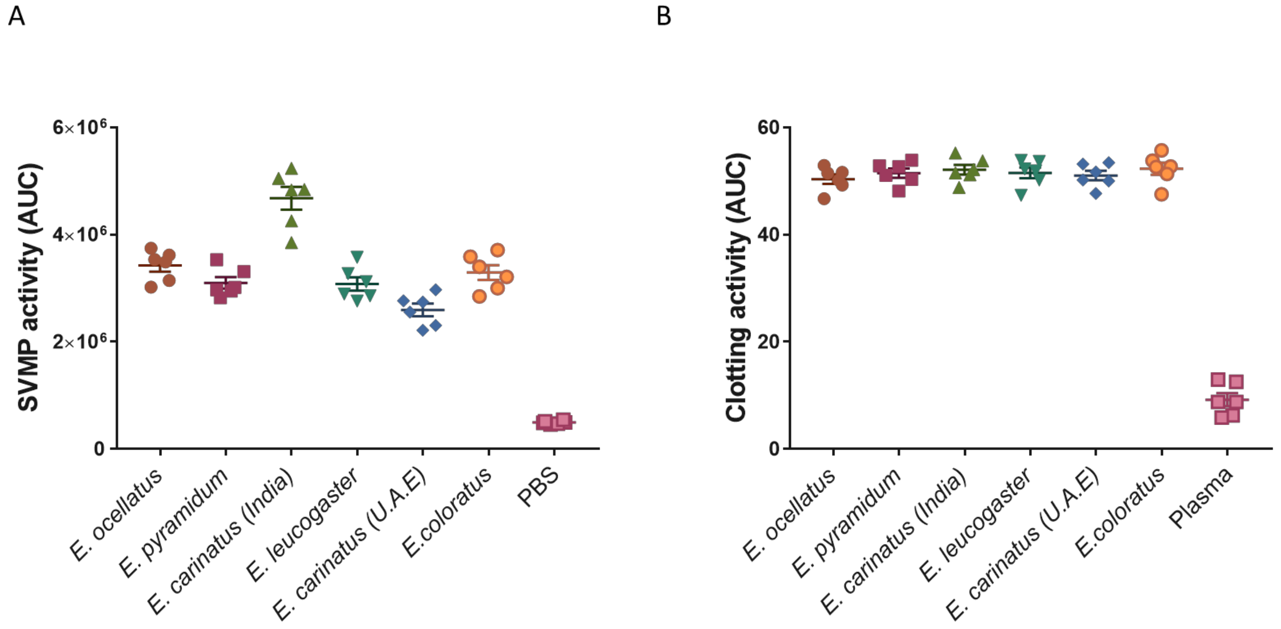


**Fig. S1. The snake venom metalloproteinase and plasma clotting activity of saw-scaled viper venoms.**

(**A**) The SVMP activity (expressed as area under the curve at 40 min) of six *Echis* venoms determined using an *in vitro* kinetic fluorescent assay. The average and SEM of n=6 repeats consisting of at least two technical replicates is represented. (**B**) The clotting activity (expressed as AUC at 60 min) of six *Echis* venoms determined using an *in vitro* kinetic absorbance assay. The average and SEM of n=6 repeats consisting of at least two technical replicates is represented.

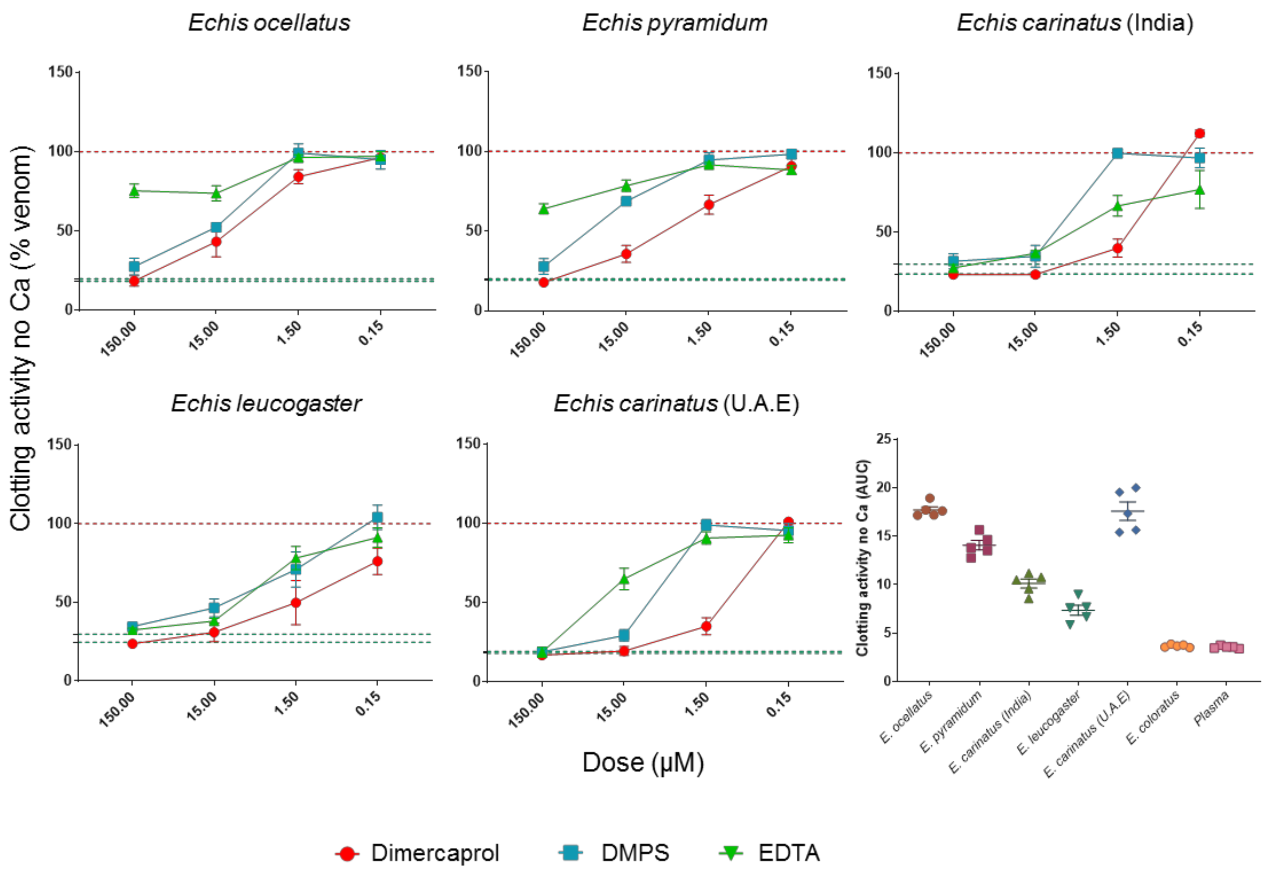


**Fig. S2. Metal chelators inhibit the procoagulant activity of saw-scaled viper venoms in the absence of calcium**.

The neutralizing capability of four metal chelators against the $Ca^{2+}$-independent procoagulant activity of six *Echis* venoms. Data is presented for four drug concentrations, from 150 µM to 150 nM (highest to lowest dose), expressed as percentages of the venom-only sample (100%, dotted red line). The negative control is presented as an interval (dotted green lines) and represents normal plasma clotting (expressed as % of venom activity). Inhibitors are color-coded (dimercaprol, red; DMPS, blue; EDTA, green). The data represents triplicate independent repeats with SEMs, where each repeat represents the average of at least two n≥2 technical replicates. Bottom right: total clotting activity of each venom expressed as area under the curve at 60 min. Note that the procoagulant activity of *E. coloratus* venom is $Ca^{2+}$-dependent, as no differences in clotting with the control are observed in the absence of calcium.

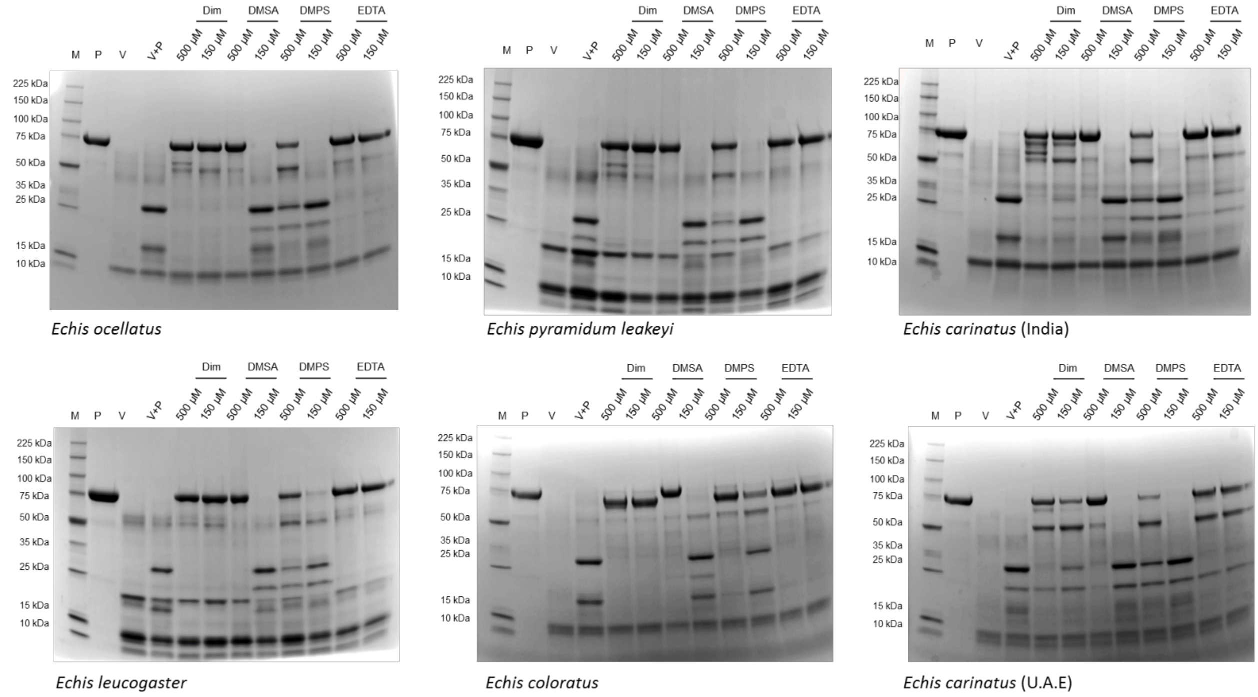


**Fig. S3. Degradation of prothrombin by saw-scaled viper venoms and inhibition of this activity by metal chelators.**

The neutralizing capability of four metal chelators against the prothrombin activating activity of six *Echis* venoms determined by SDS-PAGE gel electrophoresis. Each chelator was tested at two different concentrations (150 µM and 500 µM) and was preincubated with venom (30 min at 37 °C), followed by a 10 min incubation at 37 °C with prothrombin. Controls include prothrombin only (P), venom only (V), venom + prothrombin (V+P, positive control for degradation). M represents the protein marker.

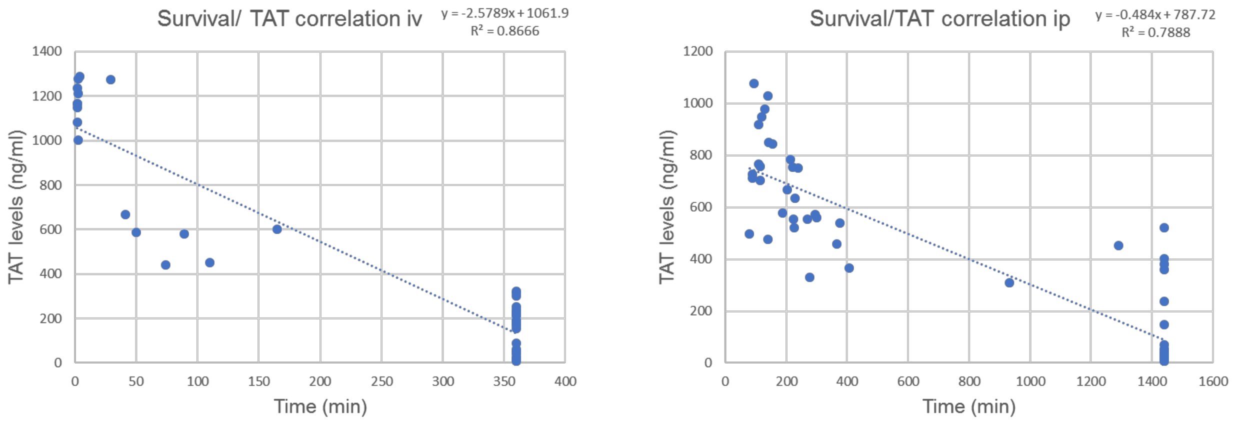


**Fig. S4. Correlations between *in vivo* survival and thrombin-antithrombin levels.**

Plots of experimental animal survival times versus thrombin-antithrombin levels. The plots represent data from animals that received intravenous treatment following a 30 min preincubation between the venom and drug (left), and those that received the intraperitoneal treatment of drug 15 mins after intraperitoneal venom delivery (right). Data points at 360 mins (left) and 1440 mins (right) represent animals that survived to the end of the experiments, and all other data points represent the time of euthanasia based on humane endpoints.

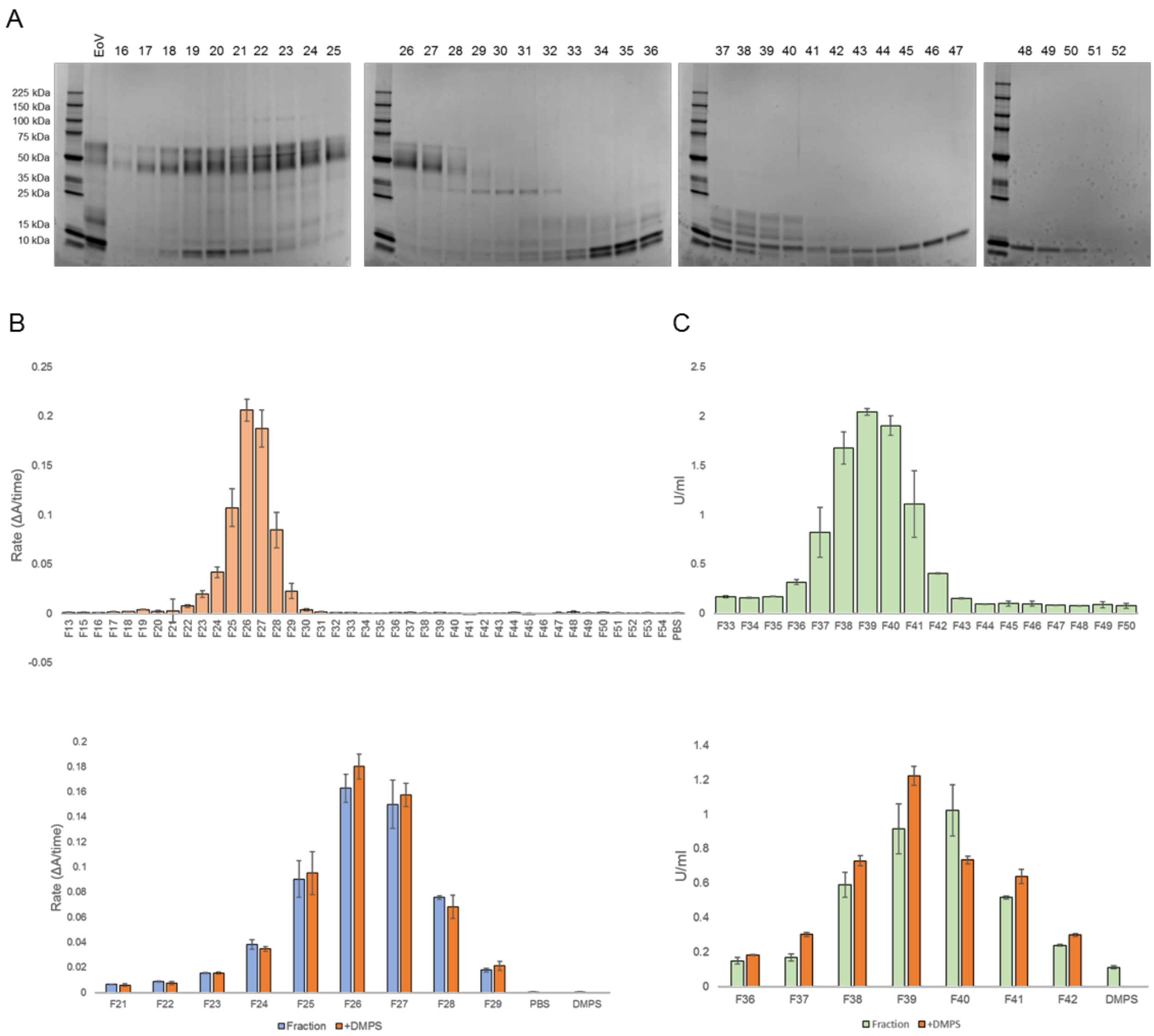


**Fig. S5. DMPS does not inhibit serine protease and PLA$_2$ activities in *E. ocellatus* venom.**

(**A**) Profiles for *E. ocellatus* venom fractions (10 µl) separated on Superdex 200 and analyzed on 4-20% SDS-PAGE gels under reducing conditions. (**B**) Serine protease activity in *E. ocellatus* venom fractions (top), and the effect of DMPS on the serine protease activity in those fractions (bottom, n=2 repeats with SDs). (**C**) PLA$_2$ activity in *E. ocellatus* venom fractions (top), and the effect of DMPS on the PLA$_2$ activity in those fractions (bottom, n=2 repeats with SDs). *The negative control has been subtracted from all samples.

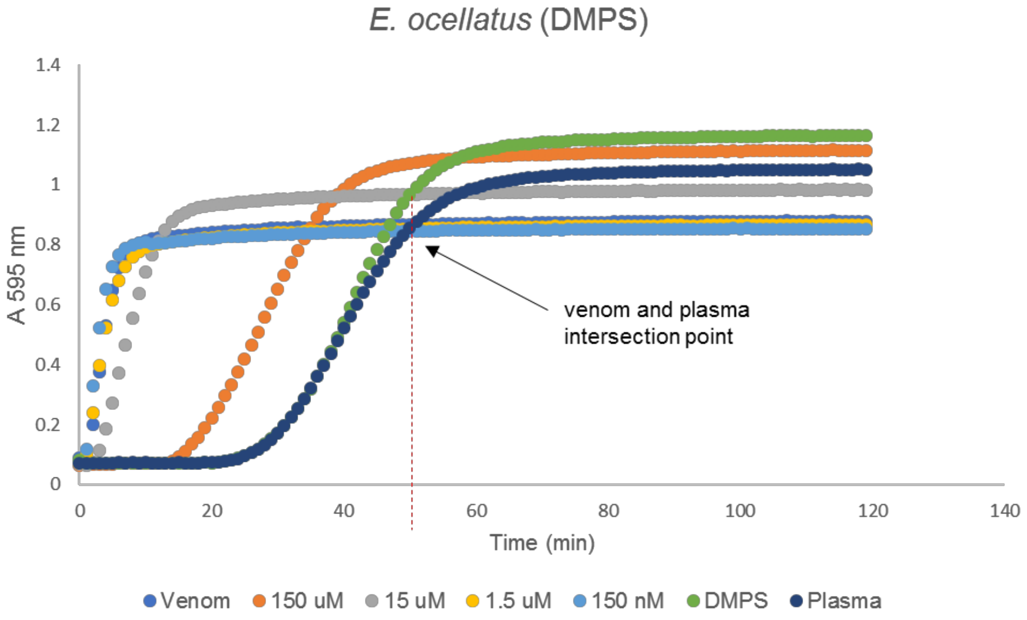


**Fig. S6. Processing of plasma clotting data.**

Plasma clotting example for *E. ocellatus* venom in the presence of DMPS. The data represents one independent repeat and the averages of triplicates within this repeat are presented. For each experiment, the intersection point of the venom and plasma samples was noted and the data up to this time point (e.g. ~50 min in this case) was considered for calculating the area under the curve. 100 ng of venom (blue) was used per assay and the concentration of the metal chelator DMPS was titrated from 150 µM to 150 nM (presented as orange, grey, yellow and light blue). The DMPS alone control was used at the highest concentration of 150 µM (green) and the plasma-only control (negative control) is shown in dark blue.
